# Chronic cold exposure induces plasticity of mitochondrial calcium uptake in beige and brown fat of UCP1-deficient mice

**DOI:** 10.64898/2026.03.16.712209

**Authors:** Casandra G. Chamorro, Sachin Pathuri, Rebeca Acin-Perez, Michael Chhan, Madeleine G. Milner, Natalia Ermolova, Anthony E. Jones, Ajit S. Divakaruni, Linsey Stiles, Andrea L. Hevener, Zhenqi Zhou, Orian S. Shirihai, Yuriy Kirichok, Ambre M. Bertholet

## Abstract

Brown adipose tissue (BAT) is a unique tissue with mitochondria specialized for thermogenesis via the BAT-specific uncoupling protein 1 (UCP1). *Ucp1^-/-^* mice cannot tolerate acute exposure to cold, illustrating the necessity of UCP1 for efficient mitochondrial thermogenesis. However, these mice adapt to low temperatures through a gradual acclimation process, suggesting a high degree of mitochondrial plasticity in brown and beige fat cells. This phenomenon, which remains to be fully elucidated, indicates the potential for these mitochondria to implement effective thermogenic mechanisms in the absence of uncoupling protein 1 (UCP1). Here, we investigated mitochondrial remodeling in beige and brown fat of *Ucp1^-/-^* mice to determine how they fulfill their thermogenic role. Upon gradual acclimation to a cold environment, Ucp*1^-/-^* mice exhibited body metabolic parameters and temperatures in the interscapular region similar to those of wild-type mice of BAT, highlighting effective thermogenesis. Interestingly, mitochondrial patch-clamp analysis and a mitochondrial Ca^2+^ swelling assay revealed a dramatic increase in Ca^2+^ uptake depending on the mitochondrial calcium uniporter (MCU) in BAT mitochondria from *Ucp1^-/-^* mice when robust thermogenesis was required. Mitochondrial remodeling was accompanied by markedly increased tethering between mitochondria and the endoplasmic reticulum (ER) in *Ucp1^-/-^*mice, confirming a significant restructuring of the contact sites between the ER and mitochondria, likely to adapt to a new Ca^2+^ homeostasis. Respiratory complexes also underwent significant reorganization, which partly led to a reduction in their assembly. Levels of ATP synthase and its F1 subcomplex increased, suggesting a major source of ATP consumption and energy expenditure. We propose a new role for MCU as a key regulator of mitochondrial plasticity, enabling efficient thermogenesis in beige and brown adipose tissues in the absence of UCP1.

## 1 INTRODUCTION

Brown adipose tissue (BAT) is a remarkable tissue with unique mitochondria that must adapt to fulfill their role in thermogenesis, depending on the environmental and body challenges encountered. To this end, these organelles possess extraordinary plasticity in their structural and functional organization, the mechanisms of which remain largely unknown. BAT therefore represents a formidable platform for discovering the different mechanisms that mitochondria might employ to preserve their thermogenic capacity. BAT is the mainstay of adaptive thermogenesis important for maintaining body temperature^1,2^ and controlling the metabolic rate^2,3^. Its recent identification in adults opens a new field of research to understand the mechanisms that control heat production to increase fat burning and energy expenditure for therapeutic purposes^4–7^. BAT activity and remodeling for optimized thermogenesis is induced under conditions that increase sympathetic tone, including prolonged exposure to cold temperatures, physical activity, or high-fat diet. This activation leads to fat burning and heat production^2–4^. Thermogenesis in brown adipocytes depends mainly on the mitochondrial uncoupling protein 1 (UCP1), carrying H^+^ across the inner mitochondrial membrane (IMM)^2,4,8^. White fat also senses these environmental changes and develops islands of adipocytes called beige adipocytes that adopt several characteristics of brown adipocytes such as UCP1 expression and multilocularity of lipid droplets^9,10^. Like brown adipocytes, beige adipocytes participate in fat burning to avoid the positive energy balance that leads to obesity and insulin resistance, and mice with these cells can resist diet-induced obesity^9–11^. Adult human adipose tissue contains a significant population of cells with a beige adipocyte molecular signature, and these cells, like BAT, can prevent obesity and type II diabetes^2,4,9–11^.

Cold therapy has been proposed for various health benefits, including cardiometabolic health linked to BAT activation^12^. In mice, exposure to cold is an excellent trigger for the plasticity of brown fat and white fat for effective thermogenesis, thereby helping maintain body temperature^13,14^. Mitochondrial thermogenesis by UCP1-dependent H^+^ transport is required for tolerance of acute cold exposure. Indeed, unlike wild-type (WT) mice, *Ucp1^-/-^* mice do not tolerate acute cold^15^. Interestingly, these mice can adapt to cold temperatures through a gradual temperature reduction^15–17^. Thus, brown and beige adipocytes develop an efficient UCP1-independent compensatory thermogenic mechanism that, at temperatures below thermoneutrality, renders *Ucp1^-/-^* mice leaner than WT mice^16,18^. Therefore, the *Ucp1^-/-^* mouse model constitutes a valuable tool for elucidating the mechanisms of beige and brown fat plasticity to support adaptative thermogenesis. Other thermogenic mechanisms beside UCP1 have been identified in beige and brown adipocytes, including the cyclic creatine/creatine-phosphate interconversion^19–22^, the ATP-driven cyclic Ca^2+^ pumping at the ER, and lipid anabolism/catabolism^23,24^. Their identification is exciting, as it potentially opens new avenues of intervention for metabolic diseases.

In this study, we discovered using the patch-clamp technique applied to mitochondria that mitochondria of brown and beige adipose tissue from *Ucp1^-/-^* mice after gradual acclimation to cold conditions show a dramatic increase in mitochondrial Ca^2+^ current via the mitochondrial calcium uniporter (MCU)^25–27^. In contrast, BAT mitochondria from WT mice develop a greater UCP1-dependent H⁺ current in response to cold challenge. The main proteins responsible for mitochondrial Ca²⁺ extrusion, NCLX and TMEM65, show decreased expression in *Ucp1^-/-^* mice compared with WT mice, suggesting a lower Ca^2+^ efflux from these transporters. However, greater ER–mitochondrial contacts (ERMCs) responsible for Ca²⁺ transfer are observed in *Ucp1^-/-^* mice. The electron transport chain (ETC) is significantly modified in *Ucp1^-/-^* mice compared to WT mice after prolonged cold exposure, with decreased activity of complexes I, II, and IV. Interestingly, complex V (ATP synthase) is increased at the protein level and also in terms of ATP hydrolysis activity. In addition, F1 subcomplexes, which are an important source of ATP hydrolysis, are significantly more numerous. An increase in Ca²⁺ uptake by mitochondria likely leads to a greater probability of mitochondrial membrane potential dissipation, which then requires complex V to work in reverse to maintain it by consuming ATP. Thus, the reverse activity of ATP synthase and the increase in the number of F1 subcomplexes may contribute to an increase in ATP consumption and, consequently, energy expenditure.

Here, we tested the hypothesis that *Ucp1^-/-^* mice undergo a drastic remodeling of their BAT mitochondria, associated with different cellular Ca^2+^ homeostasis, to support thermogenesis.

## 2 MATERIALS AND METHODS

### 2.1 Animals

Mice were maintained on a standard rodent chow diet under 12-h light and dark cycles. All animal experiments were performed with male mice (aged 2–5 months) according to procedures approved by the UCLA Institutional Animal Care and Use Committee under animal protocol ARC-2020-169 supervised by AMB. UCLA’s Animal Welfare Assurance number with the Department of Health and Human Services Office of Laboratory Animal Welfare is D16-00124 (A3196-01). UCLA’s animal care and use program is fully accredited by AAALAC, International. C57BL/6J and *Ucp1^-/-^* mice were purchased from the Jackson Laboratory. Sample sizes were based on the results of pilot experiments to ensure that statistical significance could be reached. A limitation to the generalizability of the study is that it did not consider gender/sex issues. Mice were housed at room temperature (RT) in groups with ad libitum access to food and water in a 12:12-h dark-light cycle. CL316.243 (Sigma) at 1 mg/kg was injected intraperitoneally into mice daily for 10 days. For chronic cold experiments, mice were placed individually in cages in a mouse incubator (RIS33SD RODENT INCUBATOR) for 10 days at 28°C. The temperature gradually lowered by 2°C until it reached 6°C, and the animals were kept in these conditions for 3 weeks.

### 2.2 Indirect calorimetry

Oxygen consumption, carbon dioxide production, food and water consumption, and ambulatory movement were determined in standard fed male mice using metabolic chambers (Promethion Core Metabolic System, Sable Systems International).

### 2.3. Temperature recordings

Animal colonies were maintained at 6°C. Intrascapular BAT and tail temperatures were obtained with Flir E60bx or E75 thermal cameras and analyzed with QuickReport software to extract temperature recordings of the intrascapular BAT area and tail base.

### 2.4 Histology and immunohistochemistry

Tissues (BAT, inguinal fat, epididymal fat) were dissected, fixed in 10% formalin for 24 h, and subsequently stored in 70% ethanol prior to paraffin embedding (5 mice per group). Paraffin-embedded 5-μm slices with hematoxylin/eosin (H&E) staining were produced by the Translational Pathology Core Lab at UCLA. BAT histology for lipid droplet (LD) analysis was viewed with the ImageXpress Micro XL (Molecular Devices) microscope at the Molecular Shared Screening Resource (MSSR) in the California NanoSystems Institute at UCLA. Representative H&E images were taken with Keyence BZ-X810 microscope. The In Carta software (Molecular Devices) was used for LD size analysis. For immunohistochemistry of mitochondria, slices were subjected to citrate-based antigen retrieval and then permeabilized by incubation in 0.5% Triton X-100 in phosphate-buffered saline (PBS) for 45 min. Slices were incubated for 2 h at RT with antibodies against mouse oxidative phosphorylation (OXPHOS) antibodies at a dilution of 1:50 (Abcam, ab110413). After extensive washing in PBS, secondary antibodies anti-mouse Alexa 594 (Invitrogen) at 1:500 were added for 1 h. Slides were quenched with Vector TrueVIEW Autofluorescence Quenching Kit (SP-8400-15), and coverslips were mounted with VECTASHIELD Vibrance Antifade Mounting Medium (Vector Laboratories, H-1700-10). Images were captured using an inverted microscope IX73 Olympus, X100.

### 2.5 Isolation of mitochondria and mitoplasts from BAT and subcutaneous white adipose tissue (sWAT)

Mice were euthanized by CO_2_ asphyxiation followed by cervical dislocation. For the preparation of mitoplasts, mouse BAT and sWAT tissue were isolated, rinsed, and homogenized in ice-cold medium containing 250 mM sucrose, 10 mM HEPES, 1 mM EGTA, and 0.1% bovine serum albumin (BSA) (pH adjusted to 7.2 with Trizma base) using a glass grinder with six slow strokes of a Teflon pestle rotating at 275 rotations per minute. The homogenate was centrifuged at 700*g* for 5–10 min to pellet nuclei and unbroken cells. The first nuclear pellet was resuspended in the same solution and homogenized again to increase the yield of mitochondria. Mitochondria were collected by centrifugation of the supernatant at 8,500*g* for 10 min.

Mitoplasts were produced from mitochondria using a French press. In brief, mitochondria were suspended in a solution containing 140 mM sucrose, 440 mM d-mannitol, 5 mM HEPES, and 1 mM EGTA (pH adjusted to 7.2 with Trizma base) and then subjected to a French press at 1,200–2,000 psi to rupture the outer membrane. Mitoplasts were pelleted at 10,500*g* for 15 min and resuspended for storage in 500 μl of solution containing 750 mM KCl, 100 mM HEPES, and 1 mM EGTA (pH adjusted to 7.2 with Trizma base). Mitochondria and mitoplasts were prepared at 0–4 °C and stored on ice for up to 5 h. Immediately before the electrophysiological experiments, 15–50 μl of the mitoplast suspension was added to 500 μl solution containing 150 mM KCl, 10 mM HEPES, and 1 mM EGTA (pH adjusted to 7.0 with Trizma base) and plated on 5-mm coverslips pretreated with 0.1% gelatin to reduce mitoplast adhesion.

### 2.6 Mitochondrial patch-clamp recordings

Patch-clamp recording was performed on isolated mitoplasts. These mitoplasts were 2–4 μm in diameter and typically had membrane capacitances of 0.3–1.2 pF. Gigaohm seals were formed in the bath solution containing 150 mM KCl, 10 mM HEPES, and 1 mM EGTA (pH 7 adjusted with Trizma base). Voltage steps of 250–500 mV and 1–50 ms were applied to break-in into the mitoplast and obtain the whole-mitoplast configuration, as monitored by the appearance of capacitance transients. Mitoplasts were stimulated every 5 s. Currents were normalized per membrane capacitance to obtain current densities (pA/pF).

All indicated voltages represent the voltages on the matrix side of the IMM (pipette solution) as compared to the cytosolic side (bath solution, defined to be 0 mV). Normally, currents were induced by a voltage ramp from −160 mV to +80 mV to cover all physiological voltages across the IMM. Currents flowing into mitochondria are shown as negative, and those flowing out as positive. Membrane capacitance transients observed upon application of voltage steps were removed from current traces.

Both the bath and pipette solutions were formulated to record H^+^ currents and contained only salts that dissociate into large anions and cations normally impermeant through ion channels or transporters.

Pipettes were filled with 130 mM tetramethylammonium hydroxide (TMA), 1 mM EGTA, 2 mM Tris chloride, and 100 mM HEPES. pH was adjusted to 7.5 with d-gluconic acid, and tonicity was adjusted to ∼360 mmol/kg with sucrose. Typically, pipettes had resistances of 25–35 MΩ, and the access resistance was 40–75 MΩ.

For H^+^ currents, the whole-mitoplast H^+^ current was recorded in the bath solution containing 100 mM HEPES, 150 mM sucrose, and 1 mM EGTA. For Ca^2+^ currents, the solution contained 150 mM HEPES, zero EGTA, 100 mM sucrose, and 100 µM calcium chloride. The pH was adjusted to 7.0 with Trizma base. The tonicity was ∼300 mmol/kg.

All experiments were performed under continuous perfusion of the bath solution. All electrophysiological data presented were acquired at 10 kHz and filtered at 1 kHz.

### 2.7 Immunoblot analysis

For western blot analysis, cells were lysed in radioimmunoprecipitation assay (RIPA) buffer (1% Igepal, 0.1% sodium dodecyl sulfate, 0.5% sodium deoxycholate, 150 mM NaCl, 1 mM EDTA, 50 mM Tris-HCl, pH 7.4) and a cocktail of protease inhibitors. Lysates were resolved by sodium dodecyl sulfate (SDS)–polyacrylamide gel electrophoresis (PAGE), transferred to a polyvinylidene difluoride (PVDF) membrane (Millipore), and probed with antibodies against Na^+^/K^+^-ATPase (Abcam, ab76020, GR3237646-13, 1/10,000), TOMM20 (Sigma Prestige, HPA011562, 1/500), MCU (Cell Signaling Technology, 14997, 1/100), UCP1 (Abcam, ab10983, 1/1000), NCLX (Proteintech, 21430-1-AP, 1/500), TMBIM5 (Proteintech, 16296-1-AP, 1/500), TMEM65 (Invitrogen, PA5-112762, 1/500), GRP75 (Abcam, ab53098, 1/500), Calnexin (BD Transduction Laboratories, 610523, 1/1000), VDAC (Cell Signaling Technology, 4661, 1/2000), CyD (Proteintech, 18466-1-AP, 1/1000), OXPHOS (Abcam, ab110413, 1/500), Vinculin (1:10000; V9131; Sigma-Aldrich), NDUFB6 (1:5000; #6510832 Thermo Fisher Scientific), mtCOI (1:10000; #10444035; Thermo Fisher Scientific), ATP5A (1:1,000; #439800; Thermo Fisher Scientific), SDHA (1:1,000, #459200, Thermo Fisher Scientific), SDHB (1:1,000, #ab14714, Abcam), UQCRC2 (1:1,000, #14742-1-AP; Proteintech), ATPIF1 (1:1,000, #13268S; Cell Signaling Technology), TOMM20 Alexa Fluor 488 (1:1,000; #ab205486; Abcam), and cytochrome c (1:5,000, #ab110325, Abcam).

### 2.8 Mitochondria swelling assay (BAT and liver)

BAT mitochondrial swelling assay measurements were performed according to the method of Cesura et al., 2003^28^. Swelling buffer contained 250 mM sucrose, 10mM HEPES, 2 mM potassium phosphate, 5 µM EGTA, pH 7.2. Buffer was supplemented with 5mM succinate and 2 µM rotenone as respiratory substrates. Final incubation volume was 0.2 mL and the concentration of mitochondria was 0.5 mg, where 100 µg of mitochondria were used per well. Absorbance readings were taken every 25 seconds, and the plate was shaken for 3 seconds between readings to ensure O_2_ diffusion. Data expressed as percentage changes in absorbance at 540 nm (ΔA540) versus baseline (no CaCl_2_) 210 min after the addition of 5 µM CaCl_2_.

### 2.9 Calcium uptake of BAT mitochondria

BAT mitochondrial calcium uptake measurements were performed according to Flicker et al., 2019^29^. Calcium uptake buffer (125mM sucrose, 20mM Tris-HCl, pH 7.2 with Tris base) was supplemented with 2 μM rotenone, 1mM GDP, 0.1% bovine serum albumin (fatty acid free), and 1 μM membrane impermeable Oregon Green BAPTA-6F (Thermo Fisher, O23990). Immediately prior to calcium uptake measurement, the medium was supplemented with 5mM L-glycerol-3-phosphate. A separate calcium uptake buffer was made in the same manner, except for the addition of 1µM Ruthenium Red. Fluorescence was monitored with a PerkinElmer Envision plate reader. Measurements were recorded before and after injection of 50 μM CaCl_2_ using FITC filter sets (Ex485/Em535), with a 0.5 seconds measuring interval.

### 2.10 Proximity Ligation Assay

Fixation and blocking were performed as described for immunohistochemistry. Duolink in situ detection red kit (Millipore Sigma, DUO92101-1KT) was used to verify the proximity of two antibody locations in fixed BAT samples. To label the proximity of antigens, VDAC (abcam, ab14734, 1/10,000) and IP3R (abcam, ab5804, 1/1000) antibodies were used. For each slide consisting of three tissue sections, one tissue was devoid of VDAC or IP3R respectively to detect any nonspecific binding of the Duolink PLA probes. Before mounting, tissue was quenched as described above. Slides were mounted with VECTASHIELD Vibrance Antifade Mounting Medium (Vector Laboratories, H-1700-10). Puncta were imaged at a 63x oil objective with a Leica Confocal SP8 microscope at the Advanced Light Microscopy/Spectroscopy (ALMS) Lab in the California NanoSystems Institute at UCLA. Images were analyzed using Fiji software.

### 2.11 Complex I in gel activity

BAT permeabilized mitochondria (20 μg) were run on a 3–12% native precast gel (Invitrogen). CI in gel activity was performed as previously described (Acin-Perez et al, 2020b). CI activity was stopped in 40/ methanol/10% acetic acid. Imaging was performed of the in gel activity and finally, gels were stained with Coomassie to correct for loading.

### 2.12 Complex V in gel activity

BAT permeabilized mitochondria (20 μg) were run on a 3–12% native precast gel (Invitrogen). CV in gel activity was performed as previously described (Acin-Perez et al, 2020b). CV in gel Activity was stopped in 50% methanol. Finally, gels were stained with Coomassie to correct for loading. Imaging was performed at different stages: after 3 h and O/N In Gel Activity, after fixing with 50% methanol.

### 2.13 Blue native gel electrophoresis

Mitochondria derived from BAT were permeabilized with 8 mg digitonin/mg protein. Digitonin incubation was performed on ice for 5 min and then centrifuged at 20,000*g* for 30 min as previously described (Acin-Perez et al, 2008, 2020a). Cells preparations for BN-PAGE were performed as described (Fernandez-Vizarra & Zeviani, 2021). Supernatant containing mitochondrial complexes and super complexes were mixed with Blue Native sample buffer (5% Blue G dye in 1 M 6-amiohexanoic acid), loaded, and run on a 3–12% native precast gel (Invitrogen). Gels were run till the blue front ran out of the gel and gel was transferred to PVDF membranes.

### 2.14 ATP hydrolysis from BAT extracts

ATP hydrolysis capacity or State 4 acidification rates were measured using Agilent Seahorse XF96 as described (Fernandez-del-Rio et al, 2023). Briefly, plates were loaded with 0.75–1.5 μg of mouse BAT mitochondria diluted in MAS (70 mM sucrose, 220 mM mannitol, 5 mM KH_2_PO_4_, 5 mM MgCl_2_, 1 mM EGTA, 2 mM HEPES; pH 7.2). Tissue lysates were prepared by homogenizing the tissue in Mitochondrial Assay Solution (MAS) buffer followed by a 900*g* centrifugation to remove the nuclear debris. Protein concentration was measured in the supernatant. Initial respiration of the samples was sustained by the addition of 5 mM succinate + 2 μM rotenone in the MAS after centrifugation. Injections were performed as indicated in the figure panels at the following final concentration in the well: antimycin A (AA) (2 μM), oligomycin (5 μM), FCCP (1 μM), and ATP (20 mM). To assess maximal ATP concentration, ATP was injected consecutively.

### 2.15 Determination of ATP synthesis versus hydrolysis capacity in intact mitochondria

Quantification of the ratio of hydrolysis/synthesis in BAT mitochondria fueled with either pyruvate plus malate or succinate plus rotenone (Succ+Rot) was achieved by calculating the ratio between the extracellular acidification rate (ECAR) on mitochondria started in state 4 after ATP injection (hydrolysis) and the basal oxygen consumption rate (OCR) of mitochondria started in state 3 (synthesis) before any port injection.

### 2.16 Mitochondrial content

To determine mitochondrial mass, BAT homogenate (8 μg) in 20 μl of MAS (70 mM sucrose, 220 mM mannitol, 5 mM KH_2_PO_4_, 5 mM MgCl_2_, 1 mM EGTA, 2 mM HEPES; pH 7.2) was placed in a clear-bottom 96-well microplate. Then, 130 μl of a 1:2,000 dilution of Mitotracker Deep Red FM (MTDR, ThermoFisher), a mitochondrial selective dye insensitive to weak or mild changes in membrane potential, was added and incubated for 10 min at 37°C. Plates were centrifuged at 2,000*g* for 5 min at 4°C (no brake), and supernatant was carefully removed. Finally, 100 μl PBS was added per well and MTDR fluorescence measured (λ_excitation_ = 625 nm; λ_emission_ = 670 nm). Mitochondrial content was calculated as MTDR signal (minus blank) per microgram of protein as previously described (Acin-Perez et al, 2020a).

### 2.17 Activity of complex I, II and IV in BAT homogenates

BAT homogenates, based on BCA normalization results, were set into a Seahorse XF96 microplate in 20 µl MAS buffer (2.5 and 20 µg of sample per well). For complex I activity, the medium was for NADH-induced respiration; for complex II activity, succinate + rotenone-induced respiration; and for complex IV activity, TMPD + ascorbate-induced respiration, as previously described (Acin-Perez et al, 2020a).

### 2.18 Gene expression analysis

BAT and sWAT tissues were dissected from male mice (9 per group) and snap frozen. Total RNA extraction was done using QIAzol Lysis Reagent (QIAGEN) and RNeasy Lipid Tissue Mini Kit (QIAGEN, 74804). Total RNA was reverse transcribed using a High-Capacity cDNA Reverse Transcription Kit (ThermoFisher, 4368814). Gene expression was detected in a 384-well system (Applied Biosystems, 4309849) using KAPA SYBR FAST qPCR 2x Master Mix Rox Low (Kapa Biosystems, 7959591001) and quantified on the Quant Studio 6 Flex Real-Time PCR instrument. Relative mRNA levels were normalized to expression of the housekeeping gene *RPS3*. A complete list of primers and sequences can be found below.

Mouse qRT-PCR primer sequences (gene sequence 5′ → 3′ sequence source):

RPS3 Forward, ATCAGAGAGTTGACCGCAGTTG; Reverse, AATGAACCGAAGCACACCATAG (Panic et al., 2020)

Ucp1 Forward, ACTGCCACACCTCCAGTCATT; Reverse, CTTTGCCTCACTCAGGATTGG (Kazak et al., 2015)

Dio2 Forward, CATGCTGACCTCAGAAGGGC; Reverse, CCCAGTTTAACCTGTTTGTAGGCA NCBI/Primer Blast tool

Cidea Forward, TGCTCTTCTGTATCGCCCAGT; Reverse, GCCGTGTTAAGGAATCTGCTG (Tseng et al., 2008)

Pgc1α Forward, CCCTGCCATTGTTAAGACC; Reverse, TGCTGCTGTTCCTGTTTTC (Kazak et al., 2015)

Prdm16 Forward, CCCACCAGACTTCGAGCTAC; Reverse, ATCCGTCAGCATCTCCCATC NCBI/Primer Blast tool

Phospho1 Forward, AAGCACATCATCCACAGTCCCTC; Reverse, TTGGTCTCCAGCTGTCATCCAG (Kazak et al., 2015)

Slc6a8 Forward, TGCATATCTCCAAGGTGGCAG; Reverse, CTACAAACTGGCTGTCCAGA (Kazak et al., 2015)

Gamt Forward, GCAGCCACATAAGGTTGTTCC; Reverse, CTCTTCAGACAGCGGGTACG (Kazak et al., 2015)

Gatm Forward, CAGACACAAATTGGCCGCTC; Reverse, CCCAGGTAGTTTGTAACCTGGC NCBI/Primer Blast tool

Ckmt1 Forward, GGCCTCAAAGAGGTGGAGAA; Reverse, CAGGATCTTTGGGAAGCGGT NCBI/Primer Blast tool

Ckmt2 Forward, GCATGGTGGCTGGTGATGAG; Reverse, AAACTGCCCGTGAGTAATCTTG (Kazak et al., 2015)

Serca1 Forward, GTCCGAGCAGGGCAGTATGA; Reverse, TGGTGAGAGCAGTCTCCGTG NCBI/Primer Blast tool

Serca2 Forward, TACCTGGCTATTGGCTGTTATG; Reverse, GGAAATGACTCAGCTGGTAGAA IDT/PrimerQuest Tool

Serca2b Forward, ACCTTTGCCGCTCATTTTCCAG; Reverse, AGGCTGCACACACTCTTTACC (Ikeda et al., 2017)

RyR1 Forward, ACGGAACTGTCATCAATCGCC; Reverse, TGGTGGTCGTGTTCCCAGTC NCBI/Primer Blast tool

RyR2 Forward, CCTCATTGACTCTCTGCTACAC; Reverse, GGTCTCAGTTGGCCACATATAG IDT/PrimerQuest Tool

IP3R1 Forward, GAGCAGGAGCTTGAACCAAGTC; Reverse, CCA TTGCCAAAGCTGGTAAGGC NCBI/Primer Blast tool

IP3R2 Forward, AGTGACGTCATGGCCCTGGGCCCT; Reverse, CTCTTCTGTACACACAGTATGCTT NCBI/Primer Blast tool

Cd36 Forward, TGATACTATGCCCGCCTCTCC; Reverse, TTCCCACACTCCTTTCTCCTCTAC (Oeckl et al., 2022)

Dgat1 Forward, TGGCCAGGACAGGAGTATTTTTGA; Reverse, CTCGGGCATCGTAGTTGAGCA (Oeckl et al., 2022)

Dgat2 Forward, TGCCCTACTCCAAGCCCATCACC; Reverse, TCAGTTCACCTCCAGCACCTCAGTCTC (Oeckl et al., 2022)

Lpl Forward, AGCCCCCAGTCGCCTTTCTCCT; Reverse, TGCTTTGCTGGGGTTTTCTTCATTCA (Oeckl et al., 2022)

Gk Frward, TCGTTCCAGCATTTTCAGGGTTAT; Reverse, TCAGGCATGGAGGGTTTCACTACT (Oeckl et al., 2022)

Atgl Forward, CAAGGGGTGCGCTCTGTGGATGG; Reverse, AGGCGGTAGAGATTGCGAAGGTTG (Oeckl et al., 2022)

### 2.19 Quantification and statistical analysis

Student’s t test was used to determine the significance of differences between two groups with GraphPad Prism 7. Unless otherwise specified, *p < 0.05, **p < 0.01, and ***p < 0.001. Data points and errors bars on graphs represent mean ± standard error of the mean (sem) values.

## 3 RESULTS

### 3.1 *Ucp1^-/-^* mice have functional brown fat for thermogenesis under chronic cold challenge

Because *Ucp1^-/-^* mice gradually adapt to cold despite their inability to survive acute exposure, they offer a powerful model for studying the thermogenic adaptations of BAT and beige fat. To promote mitochondrial remodeling in those tissues to induce a robust thermogenic response, we exposed mice to a cold environmental stimulus. Cold also induces the transformation of white adipose tissue into beige, a depot with high thermogenic capacity important for adaptation to chronic cold (subcutaneous fat, sWAT)^16,20,30^. We subjected mice to chronic cold adaptation for 3 weeks^16,17^ to allow *Ucp1^-/-^* mice to adapt. Muscle shivering likely contributes to the maintenance of body temperature, providing BAT and beige fat of *Ucp1^-/-^* mice time to implement alternative thermogenic mechanisms^15,16,31^. Mice were individually housed and kept for 10 days at a thermoneutral temperature of 28°C, after which the temperature was gradually lowered by 2°C until it reached 6°C, where they are maintained for 3 weeks. Body weight and food consumption were similar between the WT and *Ucp1^-/-^* groups (**Figure 1A and B**). The *Ucp1^-/-^* mice were significantly less active (**Figure 1C**), probably due to anxious behavior^32^. This, in addition to the absence of UCP1, could explain a slight but significant decrease in energy expenditure (**Figure 1D and E**). The respiratory exchange ratio (RER) was comparable between the two groups (**Figure 1F and G**). Thus, in line with previous studies, after acclimation to chronic cold exposure (6°C) for 3 weeks, most metabolic parameters in *Ucp1^-/-^* mice were comparable to those in WT mice^17^. Interestingly, BAT heat production measured using an infrared camera around the interscapular region was similar to that in the WT group (**Figure 1H and I**), suggesting robust thermogenesis in the BAT of *Ucp1^-/-^* mice, even in the absence of the main thermogenic transporter UCP1. Tail temperature also did not differ between WT and *Ucp1^-/-^* mice (**Figure 1H and I**). The thermogenic program in BAT and sWAT was essentially similar between WT and *Ucp1^-/-^* mice, suggesting that this program occurs and is therefore not dependent on UCP1 (**Supplementary Figure 1A and B**).

**Figure 1:**
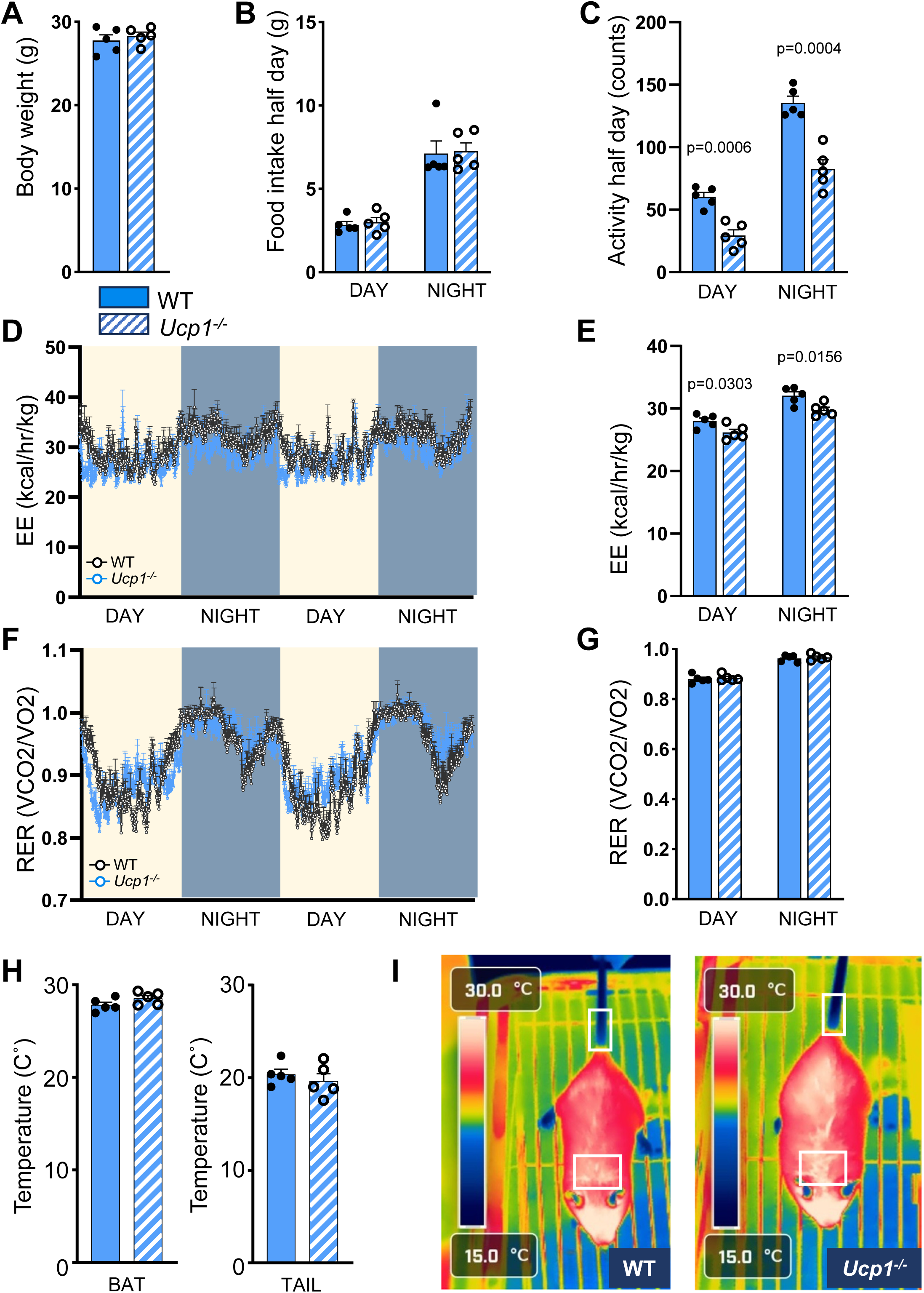
WT and *Ucp1^-/-^*mice have similar metabolic mouse phenotypes upon cold acclimation. **A to C.** Body weight (**A**), food intake (**A**), and activity (**C**) of WT and *Ucp1^-/-^* mice during the last 3 days of the 3-week cold acclimation period. **D, E.** Energy expenditure (EE). **F, G.** Respiratory exchange ratio (RER). **H.** Infrared imaging quantification of BAT and tail regions (white rectangle) of WT and *Ucp1^-/-^* mice after cold exposure. Data are expressed as mean ± sem. n = 5 per group. **I.** Representative image from the infrared camera. Unpaired t-test.

Because a considerable amount of small LDs and large mitochondrial biomass are characteristic of brown adipocytes, we assessed these two parameters, as indirect markers of BAT thermogenic activity. Interestingly, contrary to previous descriptions, H&E-stained histological sections of BAT from *Ucp1^-/-^*mice exposed to chronic cold surprisingly showed a tissue organization similar to that of WT mice, with a large population of small LDs (**Figure 2A and B, Supplementary Figure 2**). No difference in the LD number per image was observed between WT and *Ucp1^-/-^* mice (**Supplementary Figure 2C**). A small fraction of larger LDs was present in *Ucp1^-/-^* mice and absent in WT mice, but this fraction was not dominant **(Supplementary Figure 2**). A similar feature was observed in sWAT with adipocytes containing small LDs, confirming effective “beiging” of the tissue (**Supplementary Figure 3**). Thus, despite the absence of the main thermogenic mechanism implicating UCP1, *Ucp1^-/-^* mice have BAT composed largely of small LDs typical of active BAT upon chronic cold exposure. We then assessed the mitochondrial network using BAT histology. According to *in vitro* experiments on mice and human brown adipocytes, when activated, the mitochondrial network of these cells becomes fragmented to promote higher thermogenic capacity^33,34^. To visualize mitochondria directly in the BAT from mice acclimated to cold, we performed immunohistochemistry to label mitochondria with the cocktail of antibodies recognizing the OXPHOS complexes localized on the IMM and used optic microscopy to view the mitochondrial network in several brown adipocytes at once (**Figure 2C**). Mitochondria in cold-acclimated WT mice were numerous, fragmented, had a homogeneous spherical morphology throughout the tissue, and appeared to be positioned between LDs (**Figure 2C**). These were certainly cytosolic mitochondria not in direct contact with LDs, as previously described in cold-acclimated mice^35^. Mitochondria in BAT of *Ucp1^-/-^* mice also were fragmented, spherical, and located between LDs, but they appeared smaller than those of WT mice (**Figure 2C**), which could reflect different mitochondrial bioenergetics.

**Figure 2:**
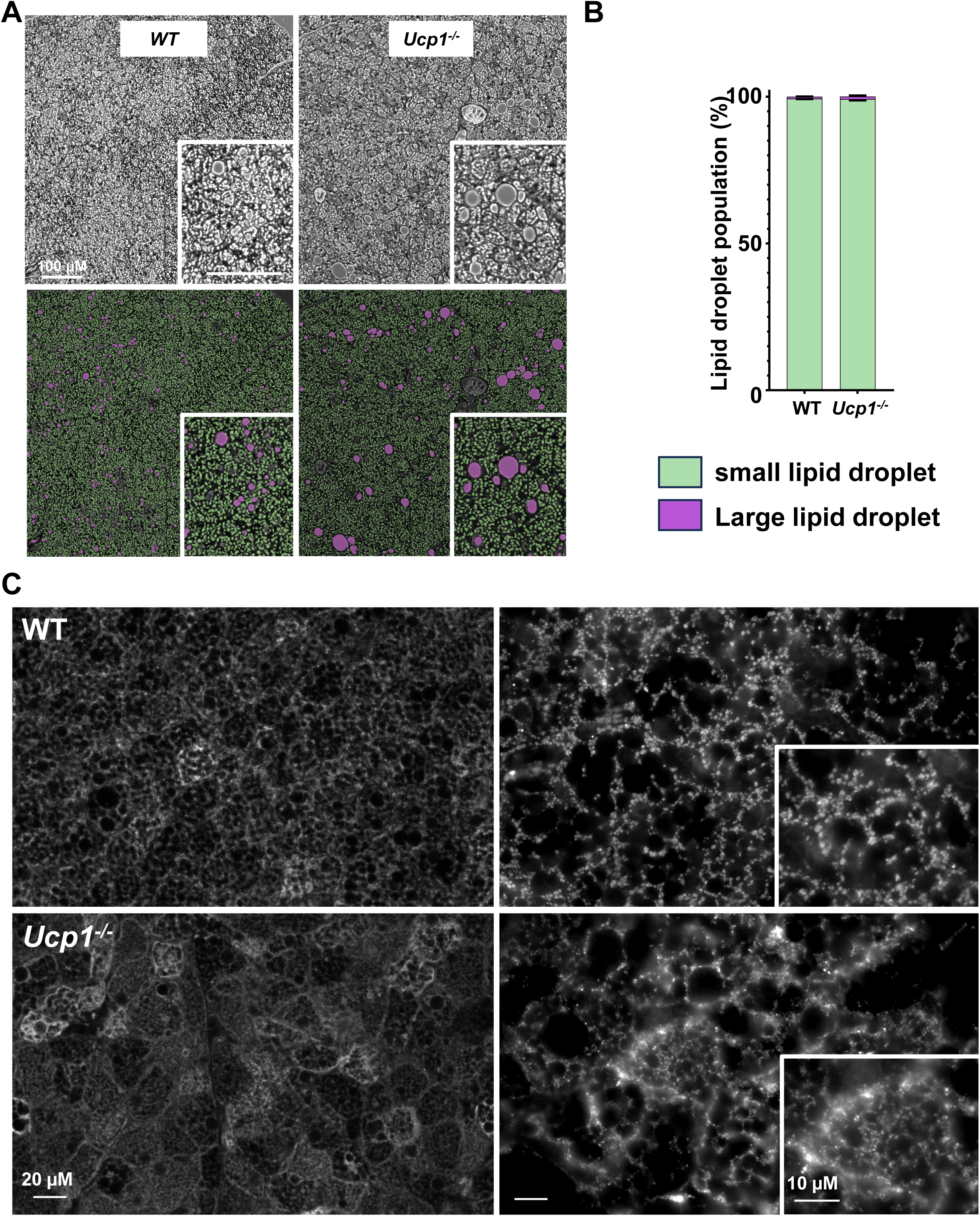
LD population and mitochondrial networks in the BAT of WT and *Ucp1^-/-^* after chronic cold adaptation are comparable. **A.** Upper panel, representative images of BAT histology after H&E staining of WT and *Ucp1^-/-^* mice upon cold acclimation (grey pictures for better quantification). Lower panel, mask built from In Carta software to evaluate LDs. **B.** LD population (small <7 µm, green; large >7 µm, purple). Data are expressed as mean ± sem. n = 4-6 mice. C. Mitochondrial morphology in BAT of WT and *Ucp1^-/-^* mice upon cold exposure was fragmented. OXPHOS immunohistochemistry. Left, magnification ×40; right, magnification ×100.

Overall, *Ucp1^-/-^* mice acclimated to a cold environment exhibited functional BAT that develops a thermogenic program, has abundant mitochondria biomass, contains numerous small LDs, and releases heat. With the absence of UCP1, a compensatory mechanism must occur for BAT to retain some thermogenic capacity with chronic cold exposure. Alternative mechanisms of UCP1-dependent thermogenesis have been previously described in *Ucp1^-/-^* BAT and sWAT and other genetic contexts^24,36^. The main proposed models involve futile cycles of ATP-consuming substrates including: 1) creatine/phosphocreatine in WT beige adipose tissue in concert with UCP1-dependent thermogenesis, 2) lipid synthesis/degradation described in *UCP1^-/-^* mice, and 3) Ca^2+^ cycling through the ER^24,36^. qPCR analysis of WT and *Ucp1^-/-^* BAT after cold adaptation showed that BAT of *Ucp1^-/-^* mice expressed higher levels of RNA transcripts for genes implicated in the creatine cycle (**Supplementary Figure 1B**) and the lipid futile cycle (**Supplementary Figure 1C**) than BAT of WT mice, suggesting a higher use of these two cycles for thermogenesis in our knockout mouse model. Ca^2+^ futile cycle genes showed similar or lower expressions, suggesting *Ucp1^-/-^* mice might use less or at least at the same level compared with WT mice (**Supplementary Figure 1D**). Overall, the three proposed futile cycles are likely to be employed at the same levels between WT and *Ucp1^-/-^* mice under chronic cold exposure.

### 3.2 The patch-clamp technique applied to mitochondria study their thermogenic ability by measuring ion fluxes across the IMM

As the mitochondria of beige and brown fats are optimized for heat production, any compensatory thermogenic mechanism can be expected to occur at the level of the IMM. The distribution of energy by mitochondria between heat and ATP depends on highly controlled fluxes of ions and metabolites across the IMM. In theory, any IMM conductance capable of dissipating the membrane potential of BAT mitochondria will result in increased ETC activity, oxygen consumption, and mitochondrial thermogenesis. Electrophysiology applied to mitochondria is the only direct method for studying those novel conductances and elucidating their thermogenic capacity (**Supplementary Figure 4**)^19,21^. We previously used the patch-clamp technique applied to mitochondria to identify novel conductances that could compensate for the loss of UCP1 and maintain their thermogenic capacity^19,21^. In this study, we first examined the UCP1-dependent H^+^ current in WT mice to reliably assess changes in H^+^ amplitude at different experimental temperatures, concomitant with different thermogenic capacities. The recording conditions were designed to measure H^+^ current across the IMM, which meant that other permeable cations were excluded from the solutions. BAT mitochondria were isolated from cold-acclimated animals and subjected to a French press to break the outer mitochondrial membrane (OMM) and expose the IMM for patch-clamp analysis (**Supplementary Figure 4**). We compared H^+^ currents in mice acclimated to thermoneutrality (28°C), when BAT is not needed to maintain body temperature, with those in mice kept at RT (mild cold) and after 3 weeks of adaptation to cold. A H^+^ current was measurable across the IMM of BAT from mice at thermoneutrality (33.53 ± 4.7 pA/pF), even when BAT mitochondrial thermogenesis was not essential (**Figure 3A and F**). The mitochondria of mice kept at RT developed a higher H^+^ current amplitude compared to those of mice kept at thermoneutrality conditions (87.63 ± 13.3 pA/pF) (**Figure 3B and F**). The largest H^+^ amplitude was recorded in BAT of mice acclimated to cold (171.61 ± 14.4 pA/pF) (**Figure 3C and F**). All H^+^ currents measured were carried by UCP1, being sensitive to GDP, a specific inhibitor of the transporter. Our previous work, in which we used β3-adrenergic agonist (CL316.243) to convert white into beige fat, showed that sWAT tissue have mitochondria that develop a UCP1-dependent H^+^ current that is two times lower than that of BAT mitochondria at RT^20^. However, during chronic cold adaptation in the current study, sWAT mitochondria exhibited a comparable H^+^ current amplitude (153.27 Pa/pF ± 19.8) to that measured in BAT mitochondria, suggesting similar thermogenic capacity when environmental challenges induce systemic changes (**Figure 3D and F**). Thus, adaptation to chronic cold provides the most favorable conditions for revealing the thermogenic capacity of BAT and sWAT mitochondria. Because BAT mitochondria primarily use H^+^ fluxes through the IMM for mitochondrial thermogenesis, we also measured whether any compensatory H^+^ current, other than the one carried by UCP1, was increased. We showed previously that ADP/ATP carrier (AAC) is the protein responsible for mitochondrial thermogenesis in non-adipose tissues^37^. AAC does not play a key role in thermogenesis in the beige and brown fat of WT mice^37^. However, as both AAC and UCP1 are proteins capable of transporting H^+^, AAC could transport H^+^ across the IMM when UCP1 is absent. The H^+^ current was not detectable in *Ucp1^-/-^* mice compared with WT mice, suggesting that AAC does not compensate for the loss of UCP1 (**Figure 3E**)^8,37,38^. Thus, *Ucp1^-/-^* mitochondria in mice exposed to chronic cold do not adapt by developing an alternative mechanism for H^+^ transport to compensate for the large UCP1-dependent H^+^ fluxes across the IMM.

**Figure 3:**
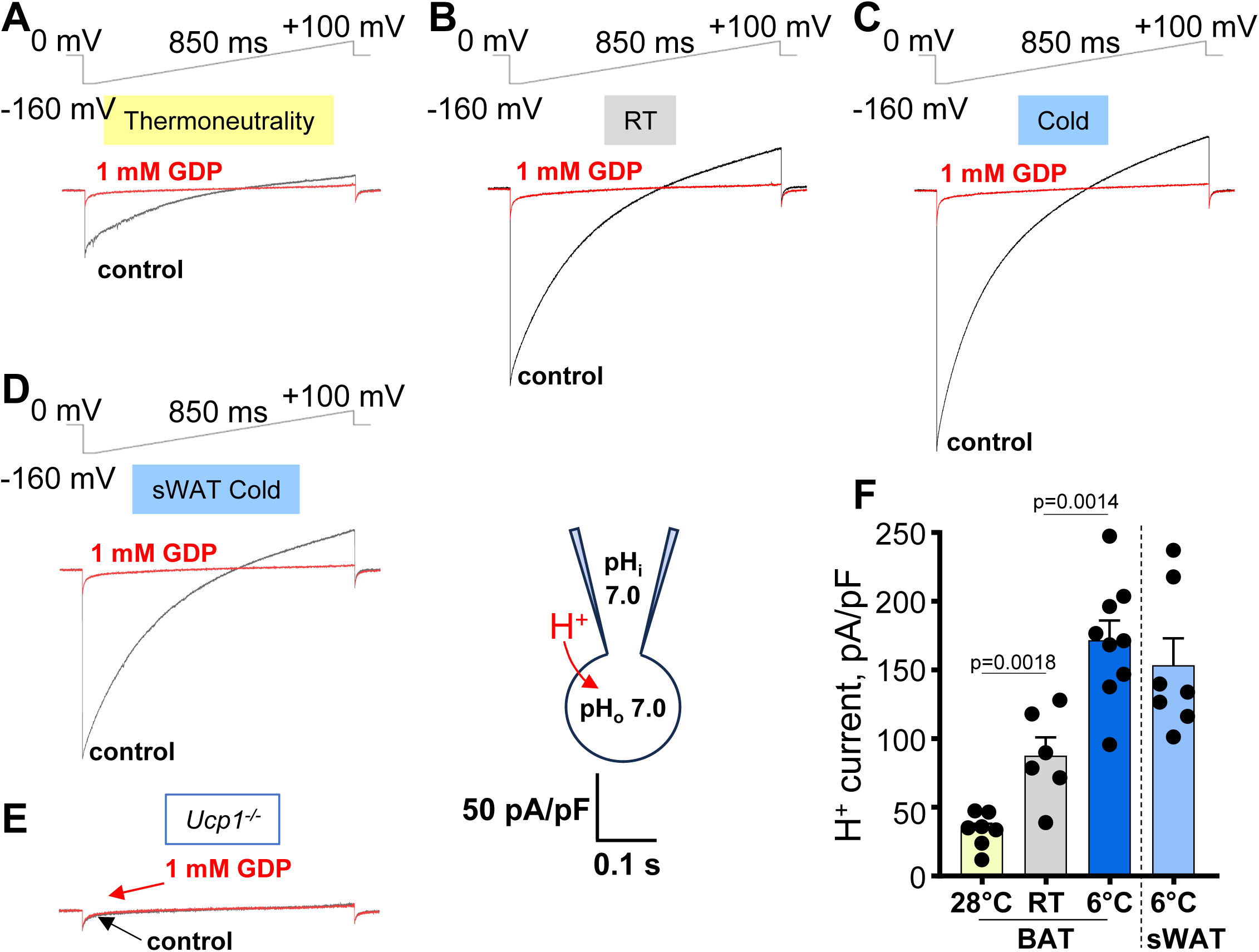
Different UCP1-dependent H^+^ current amplitudes for different environmental temperatures. **A to C.** Representative traces of the UCP1-dependent H^+^ current of BAT mitoplasts from WT mice acclimated to thermoneutrality (**A**), RT (**B**), and 6°C for 3 weeks (**C**, cold). **D.** Representative trace of UCP1-dependent H^+^ current in sWAT mitoplasts from WT mice exposed to 6°C for 3 weeks. **E.** Absence of H^+^ current in BAT mitoplasts from *Ucp1^-/-^* mice exposed to 6°C for 3 weeks. **F.** H^+^ current densities at −160 mV from **A to D**. Data are expressed as mean ± sem. n = 6–9 mitoplasts. Unpaired t-test.

### 3.3 MCU-dependent Ca^2+^ current is drastically increased in *Ucp1^-/-^* mice acclimated to chronic cold

Given that no H^+^ current was significantly upregulated, we searched for another conductance capable of dissipating the mitochondrial membrane potential and thus compensating for the loss of UCP1-dependent H^+^ current. The Ca^2+^ cation is an excellent candidate for this role, having already been proposed in other thermogenic mechanisms at the ER level^36,39^. We used recording conditions to study MCU, the main Ca^2+^ uptake mechanism for mitochondria. The MCU-dependent Ca^2+^ current was one of the first conductances characterized with the whole mitochondrial patch-clamp technique^27^. Ca^2+^ current across the IMM has a recognizable inward rectifying current^27^. Interestingly, 3 weeks of cold adaptation resulted in a dramatic increase in the Ca^2+^ current amplitude by approximately 5-fold in the BAT of *Ucp1^-/-^* mice compared with WT mice (28.5 pA/pF ± 3.7 vs 6.03 pA/pF ± 0.8, **Figure 4A and C**). At RT, the MCU-dependent Ca^2+^ current was only slightly higher in BAT mitochondria from *Ucp1^-/-^* mice (10.09 pA/pF ± 1.2) versus WT mice (6.1 pA/pF ± 0.8) (**Figure 4C**). Activation of the thermogenic program with CL316.243 for 10 days to pharmacologically activate BAT thermogenesis did not modify the Ca^2+^ current amplitude in WT (4.7 pA/pF ± 0.6) and *Ucp1^-/-^* mice compared to saline treatment (10.6 pA/pF ± 1.2 (**Figure 4B-C**). Thus, in conditions of robust thermogenic challenge like chronic cold exposure, *Ucp1^-/-^*mice develop a drastic MCU-dependent Ca^2+^ current across the IMM, which is likely not under β3-adrenergic control. Importantly, cold acclimation did not change the amplitude of the Ca^2+^ current in WT BAT mitochondria (6.03 pA/pF ± 0.8), confirming that this increase in Ca^2+^ uptake is a compensatory mechanism occurring in response to the absence of UCP1 thermogenesis (**Figure 4A-C**). Electrophysiological measurements in sWAT showed a significant two-fold increase in the Ca^2+^ current under cold conditions in *Ucp1^-/-^* mice (**Figure 4D and F**), but this Ca^2+^ current was lower than that measured in *Ucp1^-/-^* BAT (**Figure 4B and C**). β3-adrenergic agonist injection caused no difference in the Ca^2+^ currents between WT and *Ucp1^-/-^* mice sWAT mitochondria (**Figure 4E and F**). The dramatic change in the Ca^2+^ current across the IMM was confirmed by increased MCU protein levels in the BAT of *Ucp1^-/-^*mice compared to WT mice (**Figure 4G and H**), but not in sWAT mitochondria (**Supplementary Figure 5**). This increase in Ca^2+^ uptake could have positive impact on mitochondrial bioenergetics and might increase the efflux systems to regulate Ca^2+^ fluxes across the IMM^40^. The main transporters for mitochondrial Ca^2+^ extrusion are NCLX^41^, TMEM65 ^42,43^, and TMBIM5^44,45^. NCLX has been described to play a role in BAT physiology^46^. Our preliminary data showed decreases in NCLX and TMEM65 protein levels and no change in the TMBIM5 level (**Figure 4G and H**) in cold-adapted *Ucp1^-/-^* mice compared with WT mice. NCLX and TMEM65 work in concert, so a decrease in their protein levels could suggest that they are not the main pathway for our model^43^. Expression of TMBIM5, a Ca^2+^/H^+^ exchanger, was unchanged (**Figure 4G and H**). Thus, the main Ca^2+^ extrusion systems do not seem to increase Ca^2+^ fluxes in cold-adapted *Ucp1^-/-^* mice. *Ucp1^-/-^* mitochondria isolated from cold-adapted animals showed faster swelling after the addition of 50 µM Ca^2+^ compared to those from WT mice (**Figure 5A).** This swelling was based on the time needed to swell mitochondria depending on the Ca^2+^ uptake capacity. These results are consistent with the higher MCU-dependent Ca^2+^ current amplitude and higher MCU protein levels in *Ucp1^-/-^* mice after cold exposure. Although *Ucp1^-/-^* BAT mitochondria were found to swell more rapidly than those from WT mice, the swelling rate was significantly lower than that of liver mitochondria, which swell after the addition of only 5 µM Ca^2+^ (**Figure 5B**). Experiments using the Ca^2+^-impermeable probe Oregon Green 488 BAPTA-6F also showed faster Ca^2+^ uptake by BAT *Ucp1^-/-^* mitochondria compared with WT mitochondria (**Supplementary Figure 6**). Overall, these experiments confirmed the electrophysiological findings with a drastic increase in Ca^2+^ entry in mitochondria of *Ucp1^-/-^* mice acclimated to cold. *Ucp1^-/-^* BAT mitochondria can uptake more Ca^2+^ than WT mitochondria, suggesting that when UCP1 is absent, mitochondria have the capacity to uptake and process more Ca^2+^.

**Figure 4:**
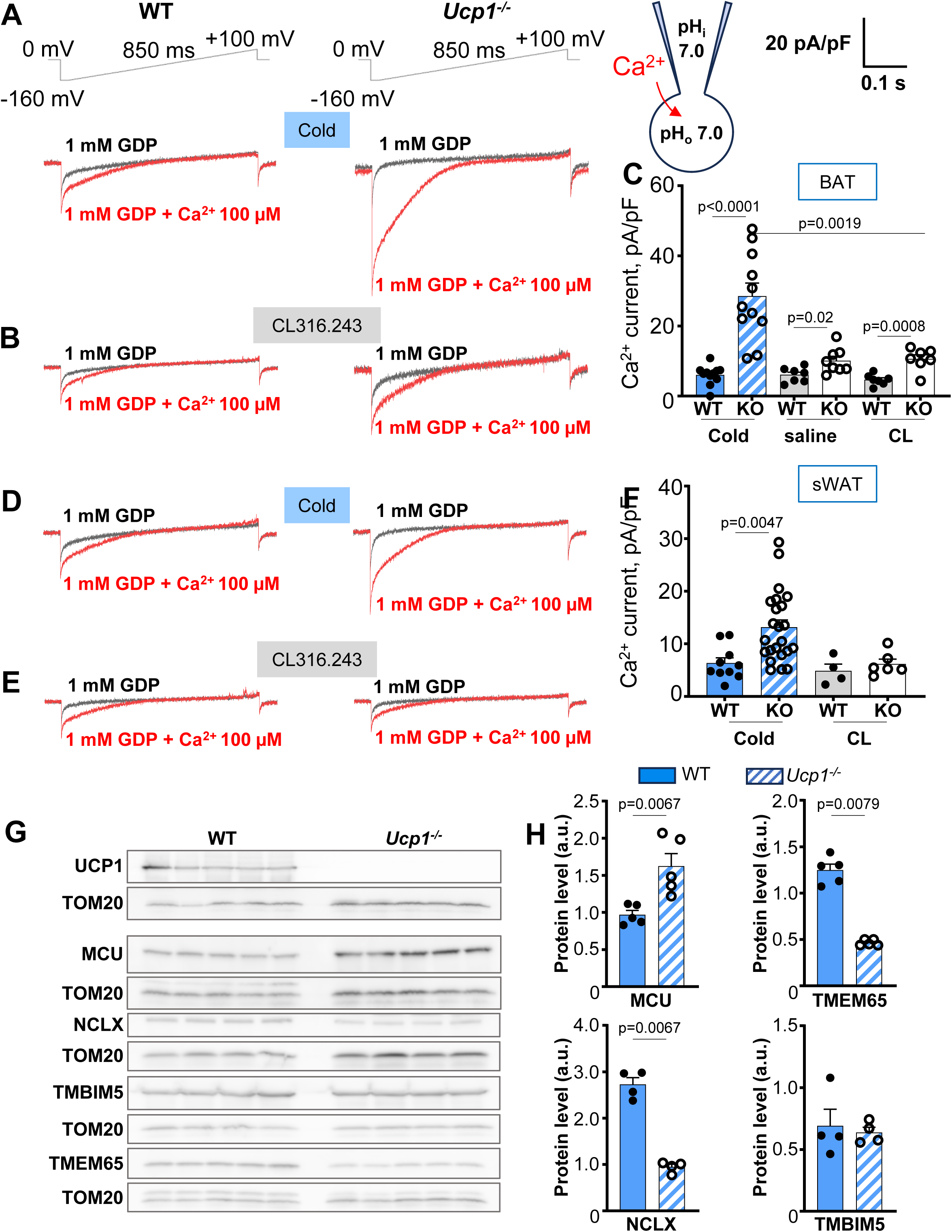
Increase in MCU-dependent Ca^2+^ currents in sWAT and BAT mitochondria of *Ucp1^-/-^* mice acclimated to chronic cold. **A.** Representative traces of MCU-dependent Ca^2+^ current in BAT mitoplasts from WT and *Ucp1^-/-^* mice exposed to 6°C for 3 weeks. **B.** Representative traces of MCU-dependent Ca^2+^ currents in BAT mitoplasts from WT and *Ucp1^-/-^* mice injected for 10 days with CL316.243. **C.** Ca^2+^ current densities at −160 mV from **A and B**. **D.** Representative traces of MCU-dependent Ca^2+^ currents in sWAT mitoplasts from WT and *Ucp1^-/-^* mice exposed to 6°C for 3 weeks. **E.** Representative traces of MCU-dependent Ca^2+^ currents in sWAT mitoplasts from WT and *Ucp1^-/-^* mice injected for 10 days with the CL316.243. F. Ca^2+^ current densities at −160 mV from **D and E**. GDP (1 mM) was added to the bath solution together with Ca^2+^ to inhibit the UCP1-dependent H^+^ current in WT mice that is present with adoption of the whole-mitoplast configuration. It allowed us to isolate MCU-dependent Ca^2+^ current from the UCP1-dependent H^+^ current. **G.** Immunoblots of proteins implicated in Ca^2+^ flux across the IMM, including MCU, NCLX, TMIM5, and TMEM65 as well as TOM20 and Na/K ATPase for normalization per mitochondrial biomass and total extract, respectively. UCP1 expression is shown to confirm the knockout profile. **H.** Protein level vs. TOM20 level under cold. N = 4-6 mice. Data shown as mean ± sem. Unpaired t-test.

**Figure 5:**
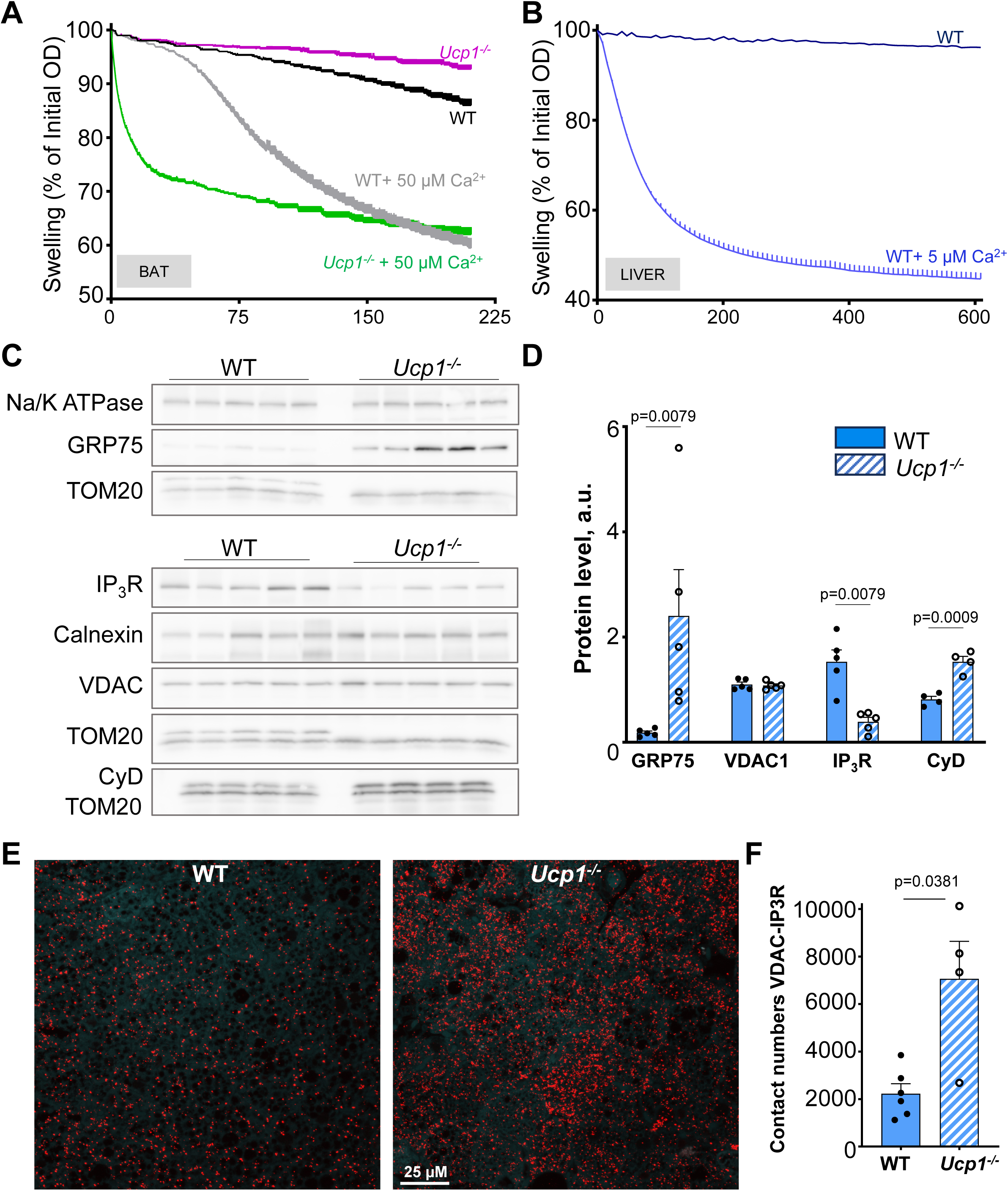
Mitochondrial calcium plasticity and ER–mitochondria contacts increase in BAT of *Ucp1^-/-^*mice upon cold exposure. A. Swelling of BAT mitochondria from WT and *Ucp1^-/-^* mice upon 3 weeks of cold exposure. **B.** Swelling of mitochondria from liver of WT mice kept at RT for comparison. **C.** Immunoblots of proteins implicated in ERMCs: GRP75, IP3R, VDAC, and cyclophilin D (CyD). **D.** Quantified protein expression. Calnexin, TOM20 and Na/K ATPase were used for normalization per ER biomass, mitochondrial biomass, and total extract, respectively. **E.** Representative image of PLA for IP3R–VDAC in BAT of WT and *Ucp1^-/-^* mice exposed to 6°C for 3 weeks. F. Quantification of the contact sites from **E**. n = 4-6 mice. Data shown as mean ± sem. Unpaired t-test.

### 3.4 ER–mitochondria tethering is increased in *Ucp1^-/-^* mice acclimated to chronic cold

Since BAT *Ucp1^-/-^* mitochondria develop a higher Ca^2+^ uptake capacity, we expect the ER network essential for Ca^2+^ homeostasis to adapt. Mitochondria are in close contact with the ER Ca^2+^ channels^47,75^. Once the cellular Ca^2+^ signaling pathway is activated, Ca^2+^ from the ER is released, leading to Ca^2+^ flux through the microdomains between mitochondria and the ER. This Ca^2+^ wave induces the MCU-dependent Ca^2+^ uptake by mitochondria^47,48^. The role of the ER in mitochondrial Ca^2+^ homeostasis is crucial, with contributions to mitochondrial functions such as ATP production, mitochondrial dynamics, metabolism, and thermogenesis^47^. The structure and functions of the interface between the ER and mitochondria are regulated and bound by several proteins located in the ER and mitochondrial membranes, which form ERMCs^49^. ERMCs are microdomains and provide a platform for metabolic and Ca^2+^ signaling^49^. One category of ERMCs specializes in Ca^2+^ flux and forms a functional complex of three proteins, playing a central role in the transfer of Ca^2+^ from the ER to the mitochondria via the MCU^50,47^. IP3 receptors (IP3Rs) are located at the ER membrane and are responsible for Ca^2+^ release. The voltage-dependent anion channel 1 (VDAC1) transports non-selective small molecules and ions such as Ca^2+^ across the OMM. Both proteins are distributed all around the membrane and at the ERMCs. The 75-kDa glucose-regulated protein (GRP75) is a mitochondrial and cytosolic chaperone that links and stabilizes the interaction between IP3R and VDAC1 to form the ERMCs^50^. This complex, the stability of which can be dynamically modulated by various stimuli, helps Ca^2+^ enter mitochondria released from the ER via IP3Rs, resulting in Ca^2+^ uptake by mitochondria.

The Ca^2+^-dependent tethering complex between the ER and mitochondria includes IP3R, VDAC, GRP75, and the recently described partner cyclophilin D^49,51^. Western blot analysis showed increased levels of GRP75 and cyclophilin D proteins in *Ucp1^-/-^* BAT compared with WT BAT under cold exposure but did not find differences in IP3R and VDAC levels (**Figure 5C and D**). To visualize and quantify the ERMCs, we performed a proximity ligation assay (PLA, **Figure 5E and F**) on BAT histology, an approach mainly used in cell lines. This is the first time this technique has been successfully applied to fixed BAT. Each punctum only forms when VDAC1 and IP3R are close enough to create a contact and for the PLA kit to add fluorophore probes, thus highlighting an ERMC. The images were obtained using a confocal microscope, and the number of puncta was analyzed using FIJI software. We found that the BAT of cold-acclimated *Ucp1^-/-^* mice had about three times more puncta than the BAT of the WT group (**Figure 5E and F**), demonstrating that mitochondria and ER form more Ca^2+^ microdomains in *Ucp1^-/-^* mice. These results confirm a significant restructuring of the contact sites between the ER and mitochondria to adapt to a new Ca^2+^ homeostasis in the absence of UCP1.

### 3.5 *Ucp1^-/-^* mice acclimated to chronic cold have fewer ETC complexes and more complex V

A recent study demonstrated a link between increased MCU protein expression and ETC dysfunction, particularly caused by respiratory complex I^52^. MCU stabilization has been proposed to support and maintain mitochondrial function when complex I is dysfunctional or diminished. Interestingly, previous studies reported a decrease in ETC complexes in the BAT of *Ucp1^-/-^* mice after cold exposure^53^. We also found that *Ucp1^-/-^* BAT exposed to chronic cold has a different OXPHOS profile. Indeed, BAT of *Ucp1^-/-^* mice showed decreases in respiratory complexes I and IV compared to WT mice (**Figure 6A and B**). The levels of complexes II and III were similar, and the only complex found at increased levels in *Ucp1^-/-^* mice compared to WT mice was ATP synthase. Respiratory supercomplexes I+III_2_, III_2_, and IV_2_ also were affected (**Figure 6C-E and G**), which corresponds to the decreases in individual complexes. *Ucp1^-/-^* mice showed an increase in the assembly level of complex V, confirming the OXPHOS profile data (**Figure 6F and G**). These structural changes in the ETC are likely to be reflected in mitochondrial respiratory activity. Oxygen consumption measurements on isolated mitochondria from WT and *Ucp1^-/-^*mice exposed to chronic cold showed an approximately two-fold decrease in the complex I-linked OCR (**Supplementary Figure 8A**). Glycerol-3-phosphate (G3P) respiration, which is standardly used for isolated BAT mitochondria respiration, also was reduced but showed a similar OCR for maximum respiratory capacity measured after addition of the chemical uncoupler FCCP (**Supplementary Figure 8B**). BAT *Ucp1^-/-^* mitochondria exhibited a coupled OCR profile, with ADP stimulating the OCR and oligomycin reducing it (**Supplementary Figure 8A and B**). Succinate/rotenone for complex II-dependent OCR (**Supplementary figure 8C**) also was slightly decreased. Respiration studies of BAT homogenates showed decreases in the CI OCR (∼6-fold), CII OCR (∼2.5-fold), and CIV OCR (∼2-fold) (**Supplementary Figure 9).** Thus, ETC activity is greatly reduced in cold-exposed *Ucp1^-/-^* mice. However, comparison of OCRs is difficult because WT mitochondria are extremely uncoupled after cold exposure. The use of GDP to inhibit UCP1 is useful, but GDP only partially blocks UCP1 uncoupling after 3 weeks of cold adaptation, which likely leads to an overestimation of differences between WT and *Ucp1^-/-^* mice. That said, the ETC content was reduced in *Ucp1^-/-^* mice, which likely results in decreased mitochondrial respiration. Interestingly, even though most of the ETC complexes decreased, complex V increases (**Figure 6A, F, and G**).

**Figure 6:**
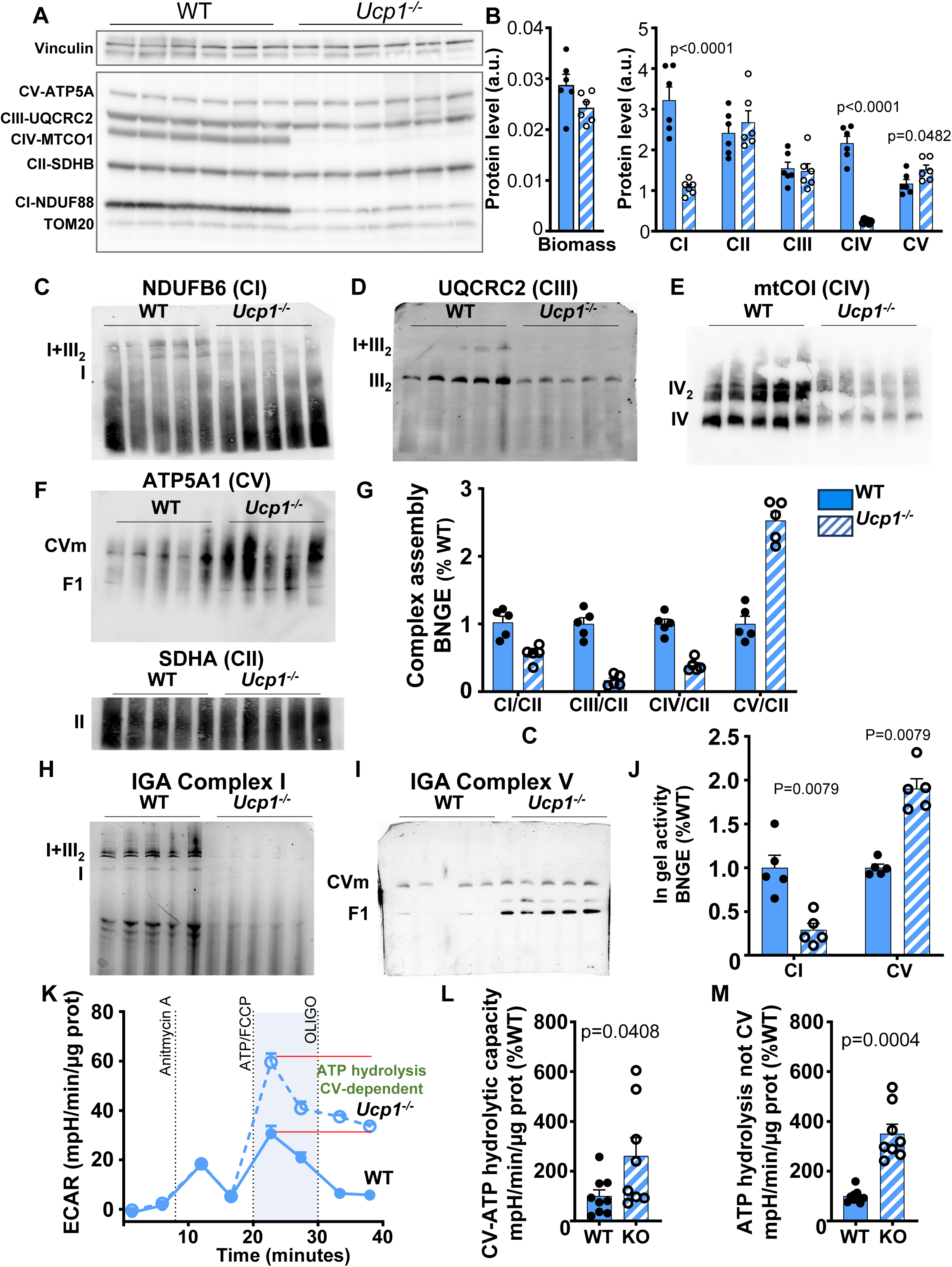
BAT of WT and *Ucp1^-/-^*mice have different ETC profiles upon cold exposure. **A.** Immunoblots showing protein levels of five mitochondrial respiratory protein complexes: NDUFB8 (complex I, CI), SDHA (complex II, CII), core 2 subunit (complex III, CIII), CIV-I subunit (complex IV, CIV), and ATP5 subunit alpha (complex V, CV) along with the loading control (TOM20) in mitochondria of BAT of WT and *Ucp1^-/-^* mice after cold exposure. **B.** Histograms of quantified protein levels based on the data presented in **A**. Data shown as mean ± sem; n = 5-6. **C to F.** Respiratory complex integrity in BAT of WT and *Ucp1^-/-^* mice upon cold exposure with I, III2, and I+III2 visualized by CI antibody (**C**), CIII antibody **(D**), complex IV (**E**), and complex V and the F1 subunit and SDHA for complex II (**F**). **G.** Complex assembly from **C** to **F**. **H and I.** In gel activity of complex I (**H**) and complex V (**I**). CVm corresponds to complex monomer (∼600 kDa). The subcomplex F1 is ∼380 KDa. **J.** Quantified in gel activity of **H** and **I**. **K.** ECAR reflecting ATP hydrolysis of complex V from BAT of WT and *Ucp1^-/-^* mice after cold exposure. **L and M.** Quantification of the ATP hydrolysis dependent on complex V (**L**) and ATP hydrolysis not coming from complex V (**M**). Data are expressed as mean ± sem. Unpaired t-test.

### 3.6 *Ucp1^-/-^* mice acclimated to chronic cold have higher mitochondrial ATP synthase activity and higher mitochondrial ATP hydrolysis activity

We next studied the activity of complex V in a native gel by ATP hydrolysis. The activity of complex I in the native gel decreased by approximately 3.5-fold (**Figure 6H and J**), confirming the reduction in complex I at the protein level. In contrast, the activity of complex V doubled (**Figure 6I and J**). Notably, three hydrolytic activities appeared on the gel (**Figure 6I**). The highest molecular weight corresponded to complex V monomers. We also found F1 subcomplexes of complex V (lowest band) and F1 subcomplexes with accessory subunits (F1-c, middle band) that were capable of hydrolytic activity. The F1 subcomplex is a matrix subunit dissociated from the F0 subunit. F1-c is an F1 subcomplex anchored to the IMM with accessory proteins but not with the F0 subunit. Both subcomplexes hydrolyze ATP but are insensitive to oligomycin, an inhibitor of complex V. We then studied the mitochondrial hydrolytic capacity of BAT homogenates using a recently reported protocol based on acidification^54^. The hydrolytic capacity of complex V (sensitive to oligomycin) in *Ucp1^-/-^* BAT was approximately 2.5 times higher than that in WT BAT (**Figure 6K and L**). ATP hydrolysis that was not inhibited by oligomycin, was 3.5 times higher in BAT of *Ucp1^-/-^* mice (**Figure 6K and M**), which is likely due to the F1 subcomplexes being dramatically increased in *Ucp1^-/-^* mice upon cold exposure. Thus, *Ucp1^-/-^* mitochondria should have more ATP hydrolysis to likely maintain the mitochondrial membrane potential due to the MCU-dependent Ca^2+^ increase. We propose that MCU-dependent Ca^2+^ entry that leads to ATP hydrolysis might increase mitochondrial energy expenditure and thermogenesis.

## 4 DISCUSSION

Mitochondrial plasticity and resilience for thermogenesis in BAT is often overlooked, and the precise molecular mechanisms that control their activation are poorly defined. This lack of information is largely due to the dearth of methods for directly measuring thermogenic proteins and other partners at the IMM level. Here, using the patch-clamp technique applied to mitochondria (**Figure 4**), we discovered that brown and beige fat mitochondria of *Ucp1^-/-^* mice, which had been progressively acclimated to cold, developed a drastic increase in mitochondrial Ca^2+^ entry via MCU^25–27^. This dramatic increase in MCU-dependent Ca^2+^ current in *Ucp1^-/-^* mice, when BAT mitochondria must provide a robust thermogenic response to cold, strongly suggests the existence of an adaptive mechanism that appears to occur only in the absence of UCP1. Because this cold-related challenge requires efficient heat production by mitochondria to maintain body temperature, we propose that MCU-mediated Ca^2+^ uptake leads to the maintenance of a mitochondrial thermogenic capacity when UCP1 is absent. Not only could this MCU-dependent adaptation mechanism be essential for *Ucp1^-/-^* mice, but it could also play a role in WT beige and brown adipocytes that do not express UCP1 but are still capable of mitochondrial thermogenesis^20,55,56^.

Consistent with previous studies, *Ucp1^-/-^* mice acclimated through 3 weeks of chronic cold exposure exhibited body metabolic parameters and interscapular heat production comparable to those of WT mice (**Figure 1**). The thermogenic gene program was also broadly comparable in the BAT and sWAT of WT and *Ucp1^-/-^* mice (**Supplementary Figure 1**). Brown adipocytes from both groups possessed similar mitochondrial biomass (**Figure 6B** and **Supplementary Figure 9D**) and multilocular LDs (**Figure 2A and B**, **Supplementary Figure 2**) consistent with active and efficient BAT under environmental cold conditions. Overall, these results suggest that *Ucp1^-/-^* mice implement a robust thermogenic program in beige and brown fat, even in the absence of UCP1, making them an excellent model for studying mitochondrial adaptation to support thermogenesis.

The role of MCU in BAT thermogenesis has been investigated recently, with one study finding that MCU does not play an essential role in BAT thermogenesis in WT mice^29^. Another study showed that MCU interacts with UCP1 to form a “thermoporter” that potentiates thermogenesis^57^, although another study revealed that MCU deficiency in mouse adipose tissue promotes energy expenditure and attenuates diet-induced obesity^58^. These different findings appear contradictory, requiring further research to characterize the role of MCU in mitochondrial thermogenesis.

Interestingly, the major mitochondrial efflux systems, NCLX and TMEM65, do not appear to increase the efflux of Ca^2+^ from the mitochondrial matrix (**Figures 4G and H**) in cold-acclimated *Ucp1^-/-^*mice. Given the very high Ca^2+^ uptake capacity of these mitochondria, it is expected that they rely on another mechanism for mitochondrial Ca^2+^ efflux. The mitochondrial Permeability Transition Pore (mPTP) has also been proposed as a Ca^2+^ efflux mechanism involved in the control of Ca^2+^ homeostasis^59^. The mPTP is a Ca^2+^-dependent channel of the IMM whose main components are AAC and ATP synthase^59^. The mPTP has different conductances, low conductance working transiently (flickering) and high conductance, operating over a long period (involved in cell death). The flickering mode could provide mitochondria with a fast Ca^2+^ release pathway for physiological Ca^2+^ homeostasis. The increased levels of ATP synthase and cyclophilin D—an activator of the mPTP (which is also part of the ERMCS complex)—suggest increased mPTP activity (**Figures 5C, 5D and 6**). We propose a Ca^2+^ cycle across the IMM, in which MCU mediates Ca^2+^ influx and the mPTP mediates its efflux.

In the present study, we found that the BAT of cold-acclimated *Ucp1^-/-^* mice had about three times more ERMCs than the WT BAT (**Figure 5E**), demonstrating that mitochondria and ER form more Ca^2+^ microdomains in *Ucp1^-/-^* mice. These results clearly confirm a significant restructuring of the contact sites between the ER and mitochondria to adapt to a new Ca^2+^ homeostasis in the absence of UCP1. These results are significant for two reasons. First, they demonstrate that *Ucp1^-/-^* brown adipocytes undergo a reorganization of key cell compartments for Ca^2+^. Secondly, the proposed model could be applicable to other cell contexts, in which mitochondrial plasticity leading to an increase in MCU requires reorganization of the ER to support a new Ca^2+^ homeostasis. We thus propose that when UCP1 is absent, a complete remodeling occurs in *Ucp1^-/-^* brown adipocytes to increase the Ca^2+^ uptake capacity to improve mitochondrial bioenergetics and thermogenesis, not only via IMM conductance but also as a key regulator of mitochondrial metabolic pathways. We therefore propose that MCU-dependent Ca^2+^ uptake, a hallmark of mitochondrial plasticity, requires a profound reorganization in brown adipocytes of *Ucp1^-/-^* mice and in brown and beige adipocytes that do not express UCP1. The β3-adrenergic pathway does not appear to be the main inducer of this plasticity (**Figure 4**), as only systemic changes such as those induced by cold exposure lead to Ca^2+^-dependent mitochondrial plasticity.

Identifying the triggers for the increase in MCU-dependent Ca^2+^ uptake will help us understand the mechanism of mitochondrial plasticity involved. The Chaudhuri laboratory has described a similar phenomenon of increased MCU-dependent Ca^2+^ uptake in models of respiratory complex I deficiency^52,60^. They concluded that the increase in MCU-dependent Ca^2+^ uptake contributes to maintaining mitochondrial energy homeostasis when mitochondrial bioenergetics is impaired^52^. They also elegantly demonstrated that MCU-dependent Ca^2+^ signaling is essential for survival during complex I dysfunction in *Drosophila*. Strengthening this mechanism, through MCU overexpression, in flies with complex I deficiency remedies both functional alterations and survival. They propose that MCU interacts directly with complex I to regulate its turnover^52^.

In the *Ucp1^-/-^* model, upon adaptation to chronic cold, we observed decreases in respiratory complexes I and IV (**Figure 6A and B**). This change in ETC composition has been described by other groups after application of a similar chronic cold protocol^17,53^. In the model of BAT developmental defects due to interleukin signaling disruption, UCP1 is also undetectable and a decrease in ETC is observed^61^. It is assumed that reductions in ETC complexes are indicative of disrupted mitochondria^53^. We believe that this assertion prevents the consideration of plasticity of mitochondria offering a remodeling mechanism dependent on CI, UCP1, and MCU. Interestingly, two studies using either a complex I dysfunction model or treatment with rotenone of CI reported a decrease in UCP1 gene expression^62,63^. We propose a functional interaction between complex I and UCP1 expression, whereby when either is decreased, mitochondrial remodeling occurs that requires MCU. Complex I is therefore part of the mitochondrial remodeling aimed at increasing Ca^2+^ uptake by mitochondria to maintain their function to support brown adipocyte thermogenesis.

Finally, the dramatic increases in the protein content of complex V and its hydrolytic activity suggest an important role for mitochondria in *Ucp1^-/-^* mice. With increased Ca^2+^ uptake, the mitochondrial membrane potential should be easily dissipated, requiring a mechanism to maintain it. The reverse activity of ATP synthase helps maintain mitochondrial membrane potential for mitochondrial bioenergetic homeostasis^54,64,65^. A recent study showed that during cold exposure, complex V has a higher hydrolysis capacity in BAT mitochondria of WT mice, which is important for BAT thermogenesis^66^. We believe that complex V functions in reverse in *Ucp1^-/-^* mice during cold adaptation to help maintain the mitochondrial membrane potential dissipated by high Ca^2+^ influx into the BAT mitochondrial matrix. We also observed a dramatic increase in F1 subcomplexes in *Ucp1^-/-^* mice, leading to increased ATP hydrolysis. Thus, the reverse-operating ATP synthase and increased F1 subcomplexes require a large pool of ATP for hydrolysis activity in BAT mitochondria of *Ucp1^-/-^* mice, which should increase energy expenditure. We propose a model in which MCU-dependent Ca^2+^ entry requires the complex V to function in reverse and more F1 subcomplexes to increase ATP hydrolysis to stimulate mitochondrial energy expenditure and thermogenesis.

Mitochondrial plasticity in *Ucp1^-/-^* mice is a remarkable phenomenon that, upon elucidation of the mechanism of induction, will offer new targets for brown adipocyte thermogenesis, with or without UCP1.

## Authorship contribution

**Casandra Chamorro**: Writing – review & editing, Visualization, Methodology, Investigation, Formal analysis, Data curation. **Sachin Pathuri, Rebecca Acin-Perez, Michael Chhan, Madeleine G. Milner, Natalia Ermolova, Anthony E. Jones**: Methodology, Investigation, Formal analysis, Data curation. Review & editing, Methodology, Data curation. **Ajit Divakaruni, Linsey Stiles, Yuriy Kirichok, Zhou Z, Hevener AL, Orian S. Shirihai:** Supervision, Funding acquisition, Project administration, Conceptualization. **Ambre M. Bertholet**: Writing – review & editing, Writing – original draft, Supervision, Funding acquisition, Project administration, Methodology, Investigation, Formal analysis, Data curation, Conceptualization.

## Acknowledgments

We thank the Bertholet laboratory for helpful discussions. We thank Drs Farha Khan and Natalia Ermolova for preliminary constructions of AAC1. We thank Israel Sekler for constructive feedback on our project. A.M.B. is supported by National Institutes of Health grants R35GM143097 and P30 DK063491, a Keck Foundation Award as well as the Pew Scholars in Biomedical Sciences. M.G.M is supported by an American Heart Association predoctoral grant 25PRE1374054. A.S.D. is supported by R35GM138003. Y.K. is supported by 2R35GM136415. Z.Z. is supported by DK125354. A.H. is supported by UCSD-UCLA Diabetes Research Center MMPC - NIH P30DK063491-23.

## Data availability

Data will be made available on request.

## SUPPLEMENTARY FIGURES

**Supplementary Figure 1: Gene expression in BAT and sWAT of WT and *Ucp1^-/-^* mice upon cold exposure. A.** Quantitative RT-PCR of canonical thermogenic genes. **B.** Quantitative RT-PCR of the main genes implicated in the creatine futile cycle. **C.** Quantitative RT-PCR of the main genes implicated in the lipid futile cycle. **D.** Quantitative RT-PCR of the main genes implicated in the calcium futile cycle. BAT profiles are in the left panel. sWAT profiles are in the right panel. Data shown as mean ± sem; n = 5 mice. Unpaired t-test.

**Supplementary Figure 2: BAT histology and LD size distribution in WT and *Ucp1^-/-^* mice upon cold exposure. A.** Representative image of H&E staining of BAT from WT and *Ucp1^-/-^* mice upon cold exposure. **B.** Relative distribution of the small LD population according to size (<7 µm in diameter). C. LD numbers per image. **D.** Relative distribution of the large LD population according to size of (>7 µm in diameter). Data shown as mean ± sem; n = 4-6 mice.

**Supplementary Figure 3: sWAT of WT and *Ucp1^-/-^* mice upon cold exposure develop beige fat profile.** Representative images of H&E-stained sWAT of WT and *Ucp1^-/-^* mice upon cold exposure.

**Supplementary Figure 4: Patch-clamp analysis of mitochondria to measure H^+^ and Ca^2+^ currents across the IMM. A.** Isolation of mitoplasts; native IMM with remnants of the OMM attached. Mitochondria were isolated from BAT and sWAT tissue lysates and exposed to a low-pressure French press to break the OMM and release the IMM, thereby generating mitoplasts. **B.** Formation of the whole mitoplast configuration of the mitochondrial patch-clamp. First, a gigaohm seal was formed between the glass patch pipette and the IMM to obtain the so-called “mitoplast-attached” configuration. Next, high-amplitude voltage pulses were applied to the pipette to break the IMM patch under the pipette (this process is called “break-in”). After break-in, the whole-mitoplast configuration formed, and the interior of the mitoplast (matrix) was perfused with the pipette solution. Since the tonicity of the pipette solution was higher than that of the bath solution, the IMM was completely released from the OMM and the mitoplast assumed a round shape. **C.** Whole-mitoplast configuration of the mitochondrial patch-clamp. Two electrodes, one in the pipette and the other in the bath, controlled the voltage across the IMM and allowed currents (I) to be recorded across the entire IMM. Convention used for the directions of currents flowing through the IMM: inward currents (positive charge entering the mitoplast) are negative, while outward currents are positive. The voltages were measured in the matrix relative to the bath solution (bath solution defined as 0 mV).

**Supplementary Figure 5: Protein profile of ERMCs in sWAT of WT and *Ucp1^-/-^* mice upon chronic cold exposure. A.** Immunoblots of proteins implicated in Ca^2+^-facilitating ERMCs: IP3R, MCU, and GRP75. **B.** Quantified protein levels. Calnexin and TOM20 were used for normalization per ER biomass and mitochondrial biomass, respectively. UCP1 data shown to confirm the knockout profile. n = 5-6 mice. Data shown as mean ± sem. Unpaired t-test.

**Supplementary Figure 6: Ca^2+^ uptake is increased in BAT mitochondria of *Ucp1^-/-^* mice upon chronic cold challenge.** Representative Ca^2+^ uptake traces of BAT mitochondria isolated from mice acclimated to chronic cold. The assay uses a non-membrane permeant Ca^2+^ probe BAPTA-6F to measure the Ca^2+^ left in the medium after uptake by mitochondria. BAT mitochondria from *Ucp1^-/-^* mice (purple trace) had a higher uptake over time than WT mice (black trace) upon injection of 50 µM Ca^2+^. This Ca^2+^ uptake is dependent of MCU, since it is inhibited by Ruthenium red (RuR).

**Supplementary Figure 7: OXPHOS profile in sWAT of WT and *Ucp1^-/-^*mice upon chronic cold exposure. A.** Immunoblots showing protein levels of five mitochondrial respiratory protein complexes: NDUFB8 (complex I, CI), SDHA (complex II, CII), core 2 subunit (complex III, CIII), CIV-I subunit (complex IV, CIV), and ATP5 subunit alpha (complex V, CV) along with the loading control (TOM20) in mitochondria of sWAT of WT and *Ucp1^-/-^* mice after cold exposure. **B.** Quantified protein levels based on the data presented in **A**. Data shown as mean ± sem; n = 5-6.

**Supplementary Figure 8: OCR in different substrates of BAT mitochondria from WT and *Ucp1^-/-^*mice upon chronic cold exposure. A.** OCR of BAT mitochondria of WT and *Ucp1^-/-^* mice acclimated to cold in pyruvate/malate respiratory substrate. **B.** OCR of BAT mitochondria of WT and *Ucp1^-/-^* mice acclimated to cold in G3P respiratory substrate. **C**. OCR of BAT mitochondria of WT and *Ucp1^-/-^* mice acclimated to cold in succinate/rotenone respiratory substrate. Data shown as mean ± sem; n = 5-6.

**Supplementary Figure 9: OCR of BAT homogenates from WT and *Ucp1^-/-^*mice after chronic cold exposure. A.** OCR of mouse BAT homogenates to measure complex II activity, (C-II) succinate + rotenone-induced respiration. **B.** OCR of mouse BAT homogenates to measure complex I and IV activities, (C-I) NADH-induced respiration, representative of complex I activity. (C-IV) TMPD + ascorbate-induced respiration, representative of complex IV activity. **C.** One-time quantification of OCR of complexes I, II, and IV in frozen BAT homogenates. **D.** Mitochondrial content. Data shown as mean ± sem; n = 5.

