## Supplementary Figures for "Chronic cold exposure induces plasticity of mitochondrial calcium uptake in beige and brown fat of UCP1-deficient mice"

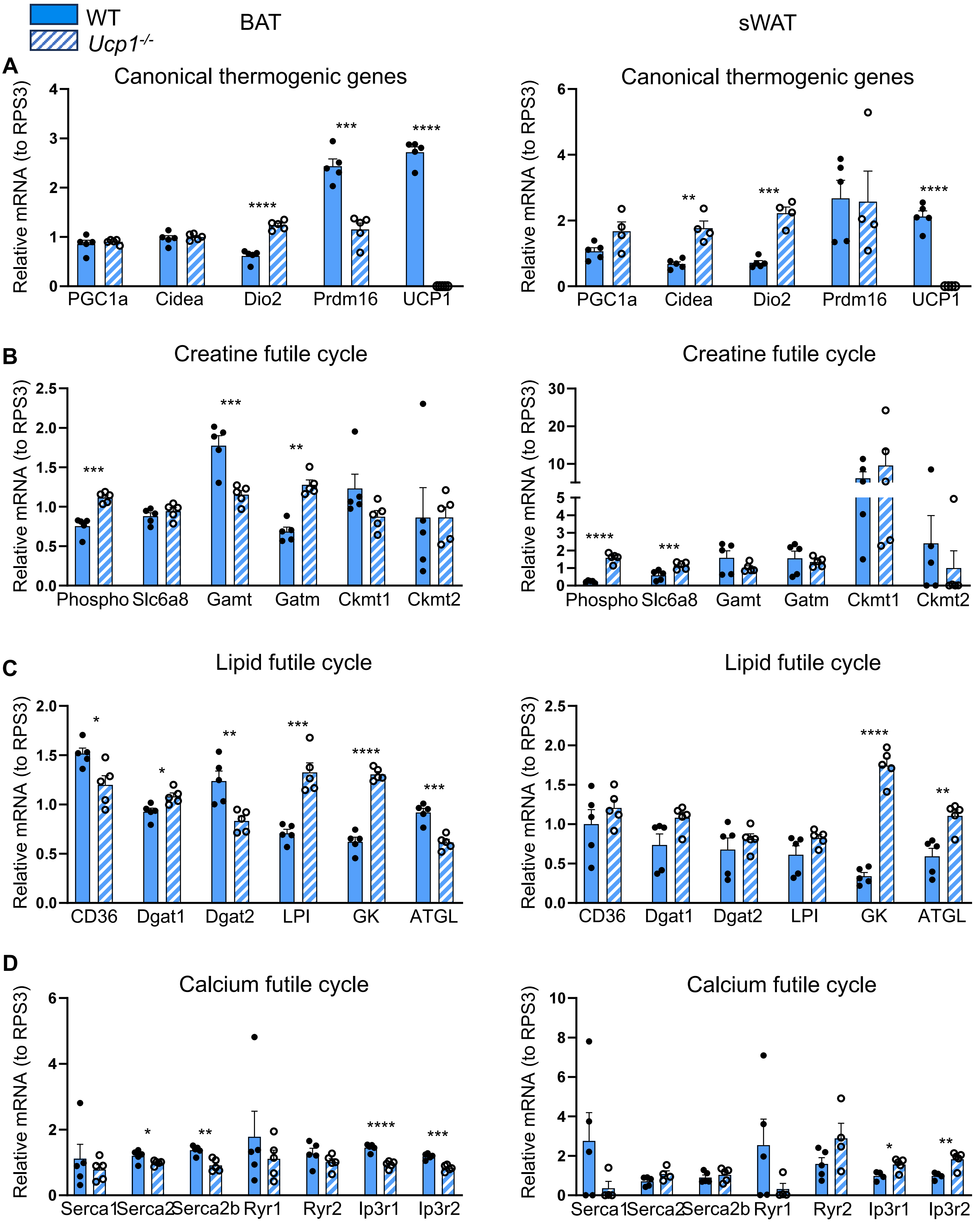


Supplementary Figure 1: gene expression of BAT and sWAT of WT and *Ucp1$^{-/-}$* mice upon cold exposure.

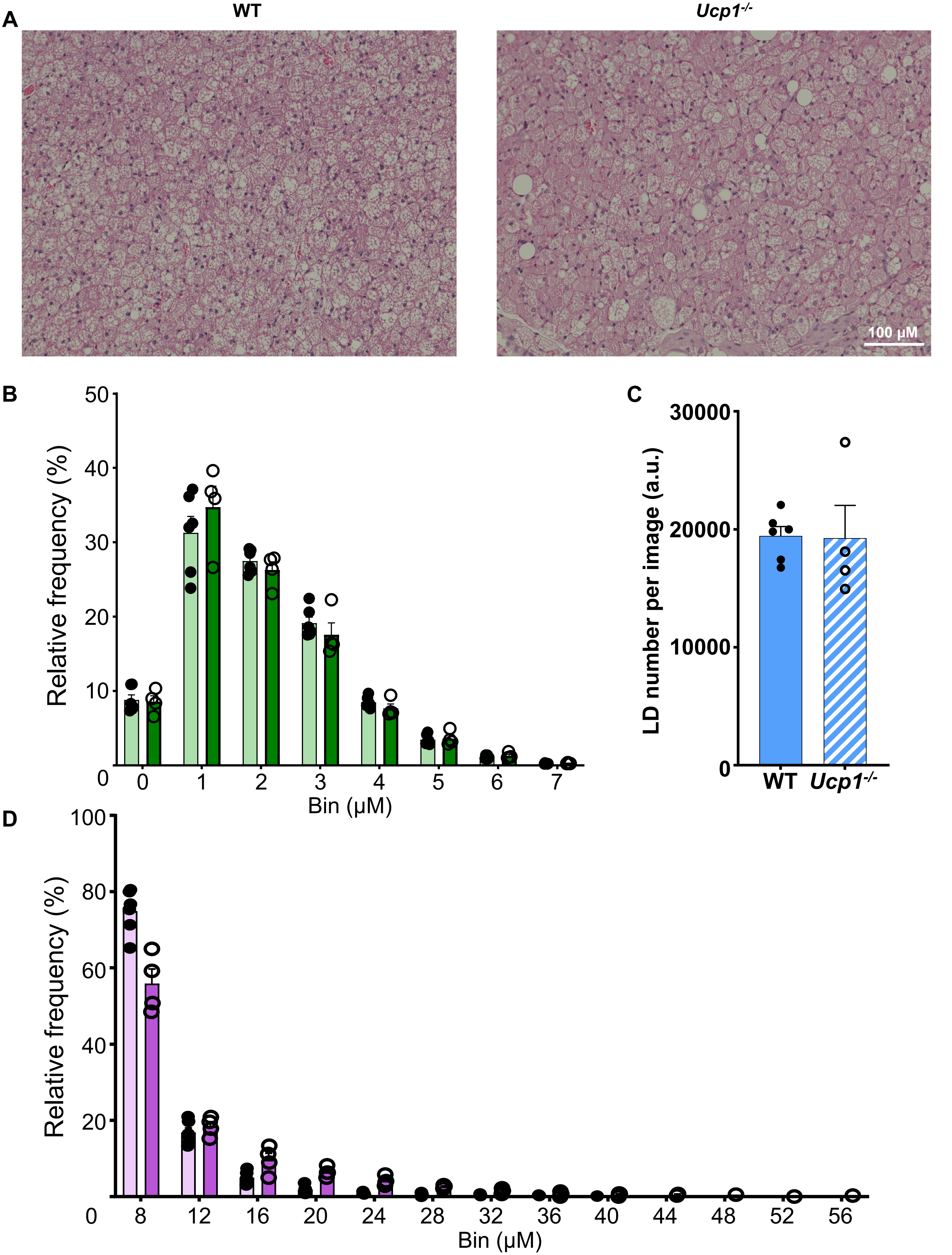


Supplementary Figure 2: BAT histology and lipid size distribution in WT and *Ucp1-/-* upon cold exposure.

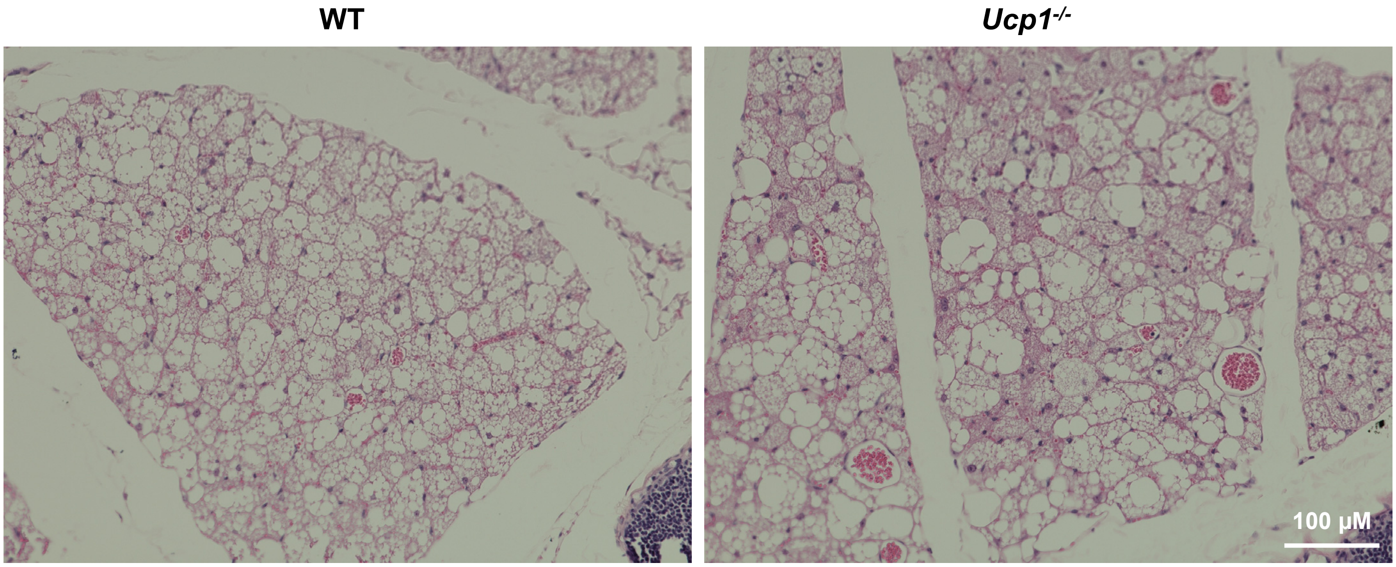


Supplementary Figure 3: sWAT of WT and *Ucp1*$^{-/-}$ upon cold exposure develop beige fat profile.

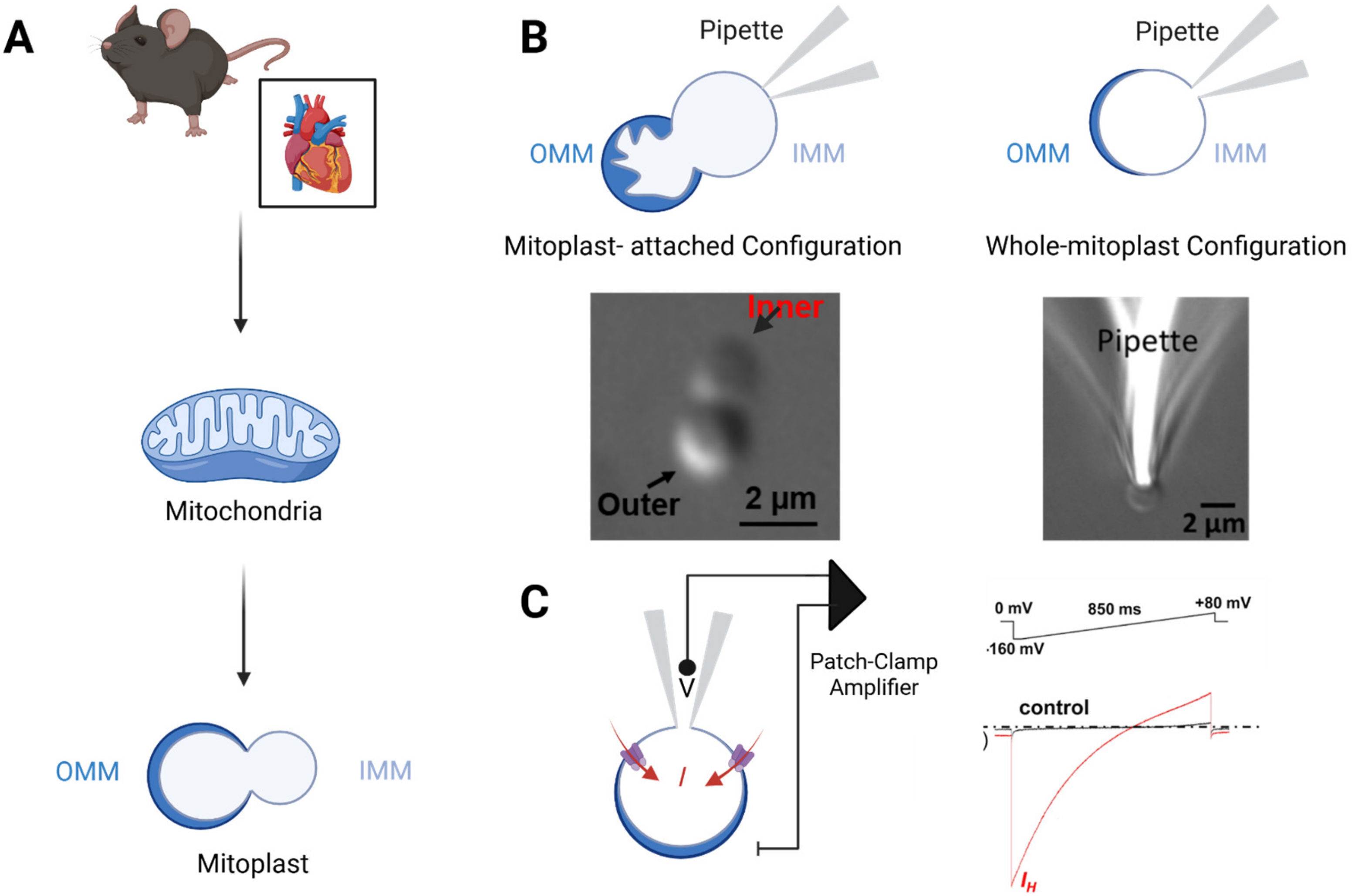


Supplementary Figure 4: Patch-clamp applied to mitochondria to measure $H^+$ and Ca2+ currents across the IMM.

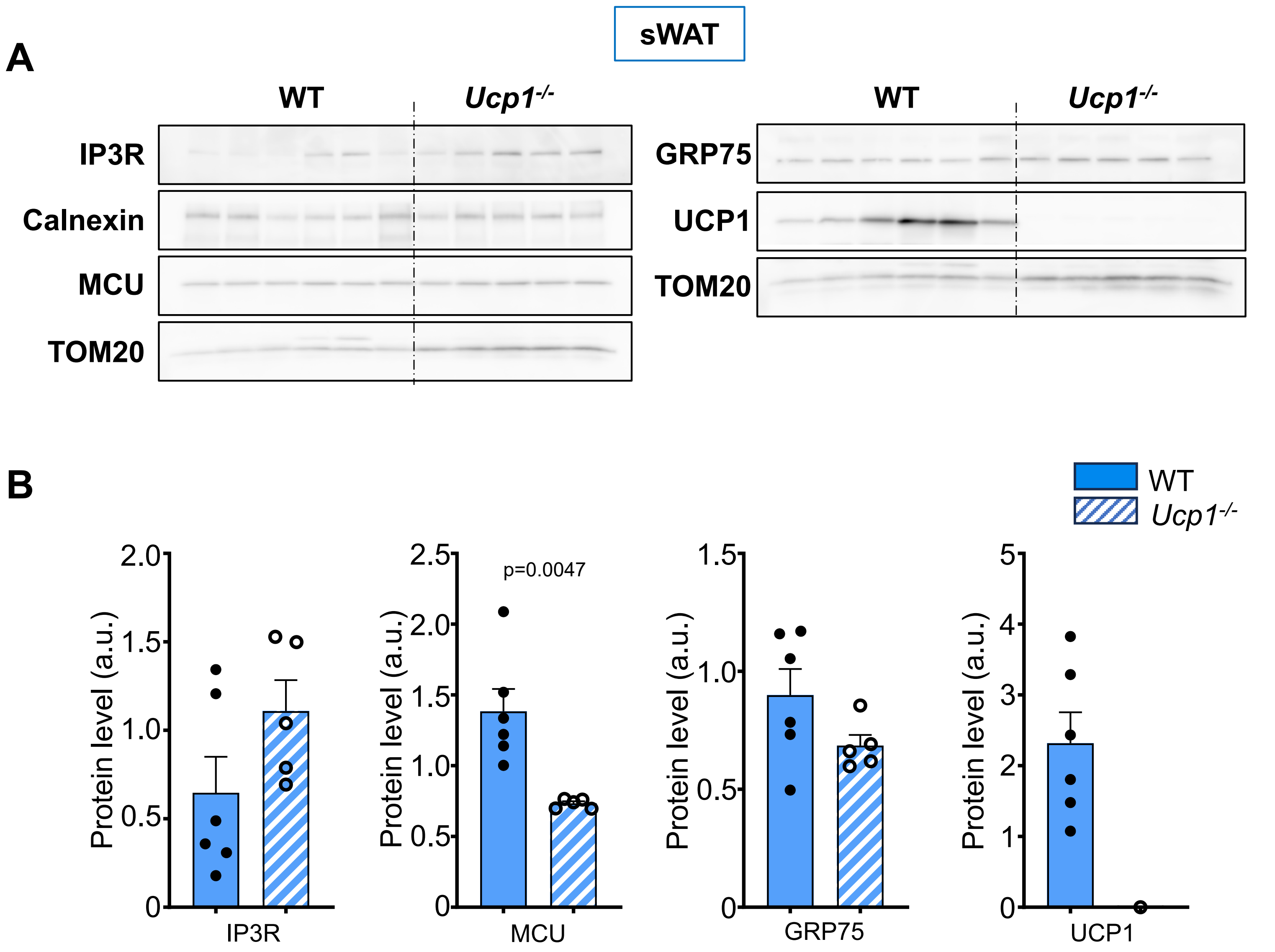


Supplementary Figure 5: Protein profile of ERMCS in sWAT of WT and *Ucp1*$^{-/-}$ mice upon chronic cold exposure.

A

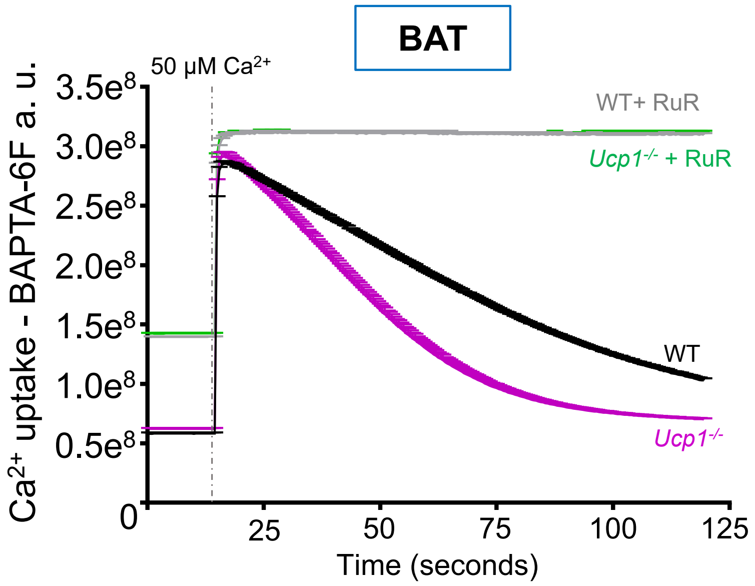


Supplementary Figure 6: $Ca^{2+}$ uptake in is increased in BAT mitochondria of *Ucp1*$^{-/-}$ mice when challenged to cold environment.

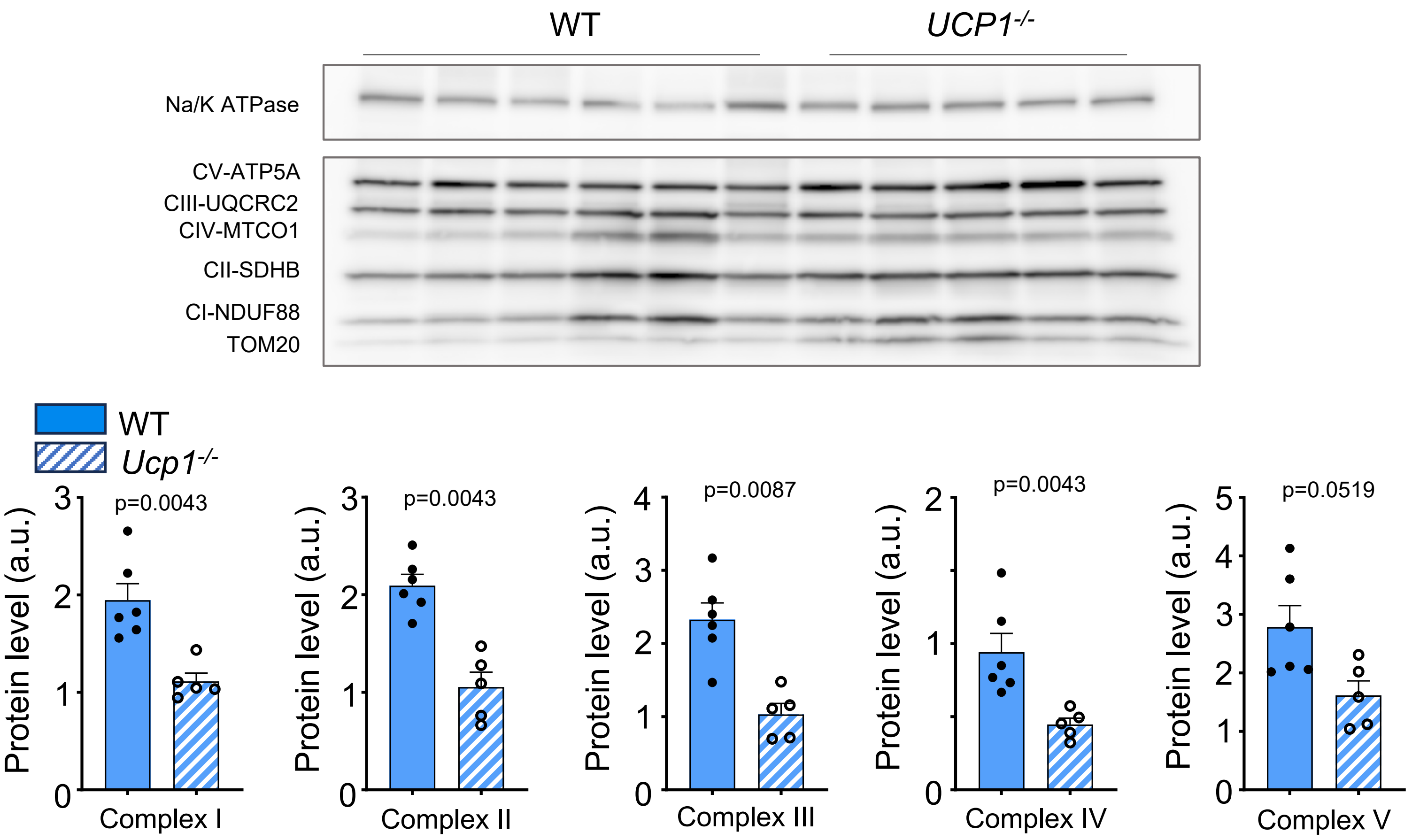


Supplementary Figure 7: OXPHOS profile in sWAT of WT and *Ucp1*$^{-/-}$ mice upon chronic cold exposure

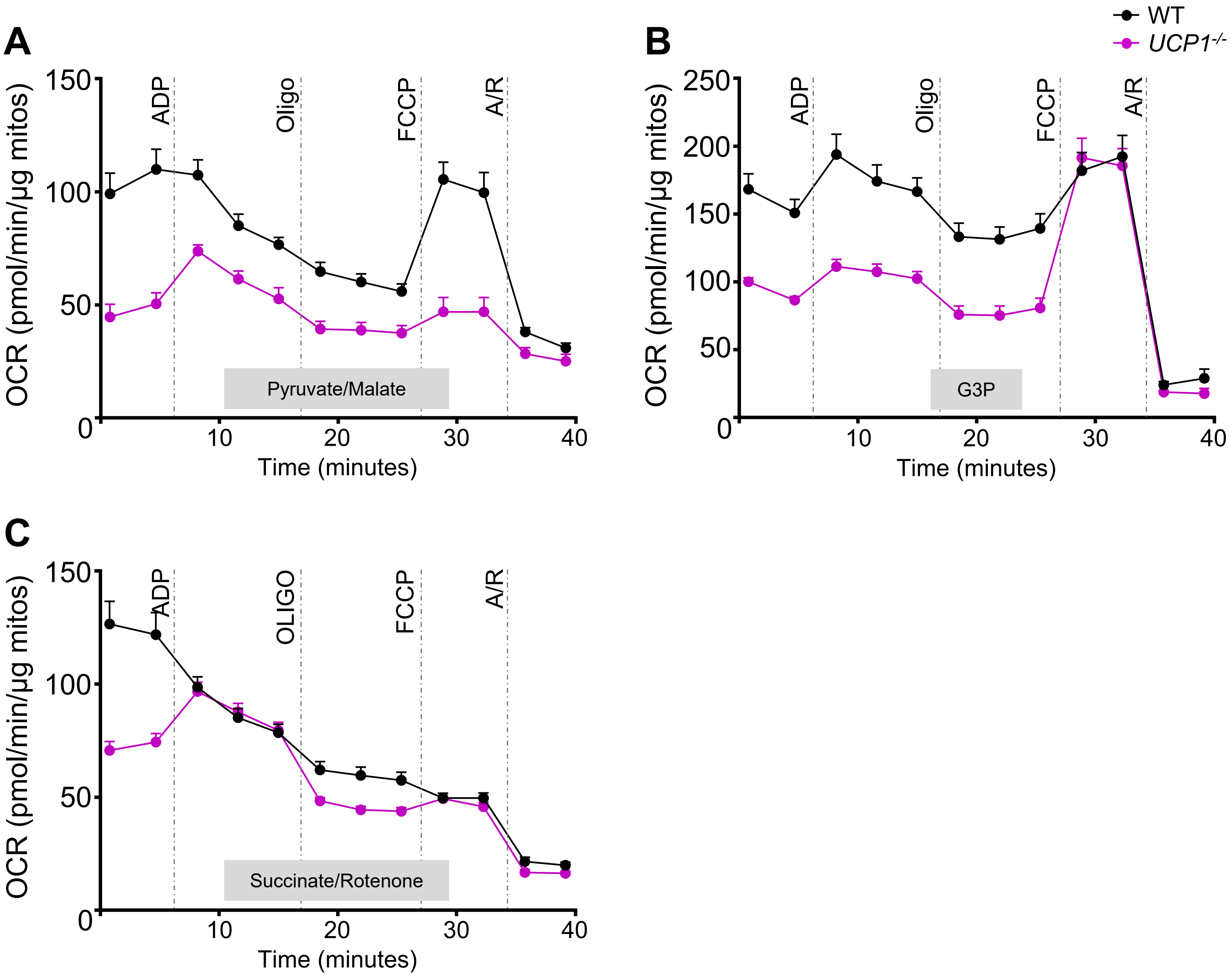


Supplementary Figure 8: OCR of BAT mitochondria in different substrates of WT and *Ucp1*$^{-/-}$ mice upon chronic cold exposure.

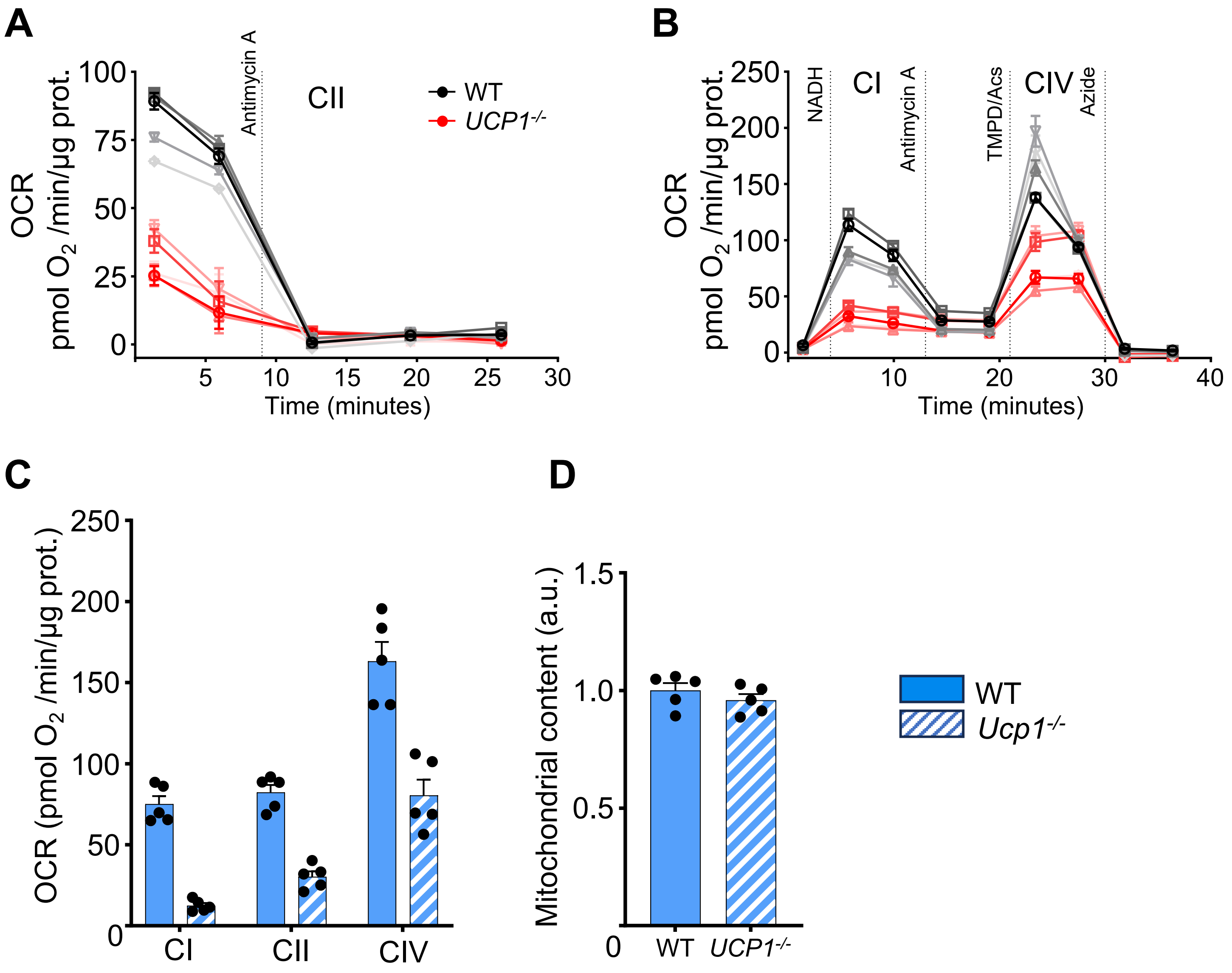


Supplementary Figure 9: OCR of BAT homogenates of WT and *Ucp1*$^{-/-}$ mice after chronic cold.
